# NlpE is an OmpA-associated outer membrane sensor of the Cpx envelope stress response

**DOI:** 10.1101/2022.10.18.512811

**Authors:** Timothy H. S. Cho, Junshu Wang, Tracy L. Raivio

## Abstract

Gram-negative bacteria utilize several envelope stress responses (ESRs) to sense and respond to diverse signals within a multi-layered cell envelope. The CpxRA ESR responds to multiple stresses that perturb envelope protein homeostasis. Signaling in the Cpx response is regulated by auxiliary factors such as the outer membrane (OM) lipoprotein NlpE, an activator of the response. NlpE communicates adhesion to surfaces to the Cpx response; however, the mechanism by which NlpE accomplishes this remains unknown. In this study, we report a novel interaction between NlpE and the abundant OM protein OmpA. Both NlpE and OmpA are required to activate the Cpx response in surface-adhered cells. Furthermore, NlpE senses OmpA overexpression and the NlpE C-terminal domain transduces this signal to the Cpx response, revealing a novel signaling function for this domain. Overall, these findings reveal NlpE to be a versatile envelope sensor that takes advantage of its structure, localization, and cooperation with other envelope proteins to initiate adaptation to diverse signals.

**Significance:** The envelope is not only a barrier that protects bacteria from the environment but also a crucial site for the transduction of signals critical for colonization and pathogenesis. The discovery of novel complexes between NlpE and OmpA contributes to an emerging understanding of the key contribution of complexes of β-barrel proteins and lipoproteins to envelope stress signaling. Overall, our findings provide mechanistic insight into how the Cpx response senses signals relevant to surface adhesion and biofilm growth to facilitate bacterial adaptation.

## Introduction

Gram-negative bacteria such as *Escherichia coli* possess dozens of two-component signal transduction pathways which regulate diverse and key cellular processes (1). Often, sensor kinases are inner membrane (IM)-bound; thus, signals originating outside of the cell or in the envelope itself must be transduced through the envelope in order to activate these systems. However, signaling in and across the Gram-negative envelope is complicated by several unique characteristics. The periplasmic space between the IM and outer membrane (OM) lacks ATP, an essential factor in phosphorylation events commonly used in signal transduction. The envelope is also a spatially complex organelle where the IM and OM are physically separated by a peptidoglycan cell wall, and signals in distinct compartments of the envelope must also navigate through this layer in order to activate IM sensors.

The Cpx envelope stress response is a major adaptive system of Gram-negative bacteria that responds to stresses that disrupt envelope protein folding and homeostasis, particularly at the IM (2). As a canonical two-component system, the Cpx response utilizes an IM sensor kinase, CpxA, to phosphorylate and modulate the activity of a DNA-binding transcriptional regulator, CpxR. Signaling by the Cpx response is also regulated by at least two other factors: CpxP and NlpE. The periplasmic chaperone-like protein CpxP inhibits activation of the system in the absence of specific cues (3, 4). In contrast, the OM lipoprotein NlpE is an activator of the response. Early studies of the Cpx response identified NlpE as an activator of the Cpx pathway when overproduced (5), and subsequent studies suggest that NlpE also senses cell-surface events, namely, adhesion to hydrophobic surfaces and certain epithelial cell lines (6, 7). The most recent studies of NlpE suggest that NlpE functions as a sensor for defects in OM lipoprotein biogenesis (8-10); chemical agents or mutations that disrupt lipoprotein processing and trafficking disrupt NlpE biogenesis, allowing NlpE to signal the presence of these stresses to the Cpx response, presumably due to its mislocalization to the IM. New NlpE synthesis is required to sense impaired lipoprotein biogenesis, suggesting that NlpE continuously functions as a cellular indicator of OM lipoprotein health (9).

NlpE’s structure was solved as a domain-swapped dimer with each monomer consisting of independently-folding N- and C-terminal domains connected by a linker region (11). This structure lends itself to a model for signaling from the OM where OM-anchored NlpE uses the C-terminal domain to reach across the periplasm to activate CpxA under certain inducing cues (e.g. surface adhesion). In the hypothetical monomer structure proposed, secondary structure elements largely limit the length of the region between the N- and C-terminal domains when NlpE is fully-folded, leading Hirano and colleagues to speculate that selective unfolding of the N-terminal domain may extend the C-terminal domain into the periplasm to facilitate activation of CpxA. Current studies have established that the N-terminal domain is sufficient for activation of the response when NlpE is overexpressed or mislocalized to the IM, suggesting that NlpE-CpxA interactions at the IM occur independently of the C-terminal domain (9, 10). Delhaye and colleagues have shown the disulfide bond in the C-terminal domain of NlpE may be involved in sensing periplasmic redox state, although it is unclear how this is communicated to CpxA (10). This study is the only one we are aware of which has provided experimental evidence for the function of the C-terminal domain. Significantly, no studies have yet elucidated how NlpE might signal from the OM, where the NTD is presumably localized in close proximity to the OM by virtue of the acylated N-terminus (11).

In this study, we investigate the mechanisms of NlpE signaling from its OM context. We report a novel interaction between NlpE and the major OM protein OmpA. Both NlpE and OmpA are required to activate the Cpx response in surface adhered cells. Further, NlpE is also sensitive to the level of OmpA in the envelope and signals OmpA overexpression to the Cpx response via its C-terminal domain. Taken together, these results demonstrate that NlpE is a bona fide OM sensor that uses its localization, multidomain structure, and interactions with other envelope proteins to sense and transduce diverse signals to the Cpx response.

## Results

### NlpE interacts with OmpA

We began this study by generating a set of expression vectors expressing C-terminally His-tagged NlpE. These vectors expressed a series of NlpE truncations from both the C- and N-terminus while preserving the native signal sequence, lipobox, and localization signals (Figure 1AB). Western blot analysis showed that variants containing an intact N-terminal domain are stably expressed, while those lacking a complete N-terminal domain showed decreased stability (Figure S1A). To study NlpE protein-protein interactions *in vivo*, we conducted disuccinimidyl suberate (DSS) crosslinking on strains expressing these variants (Figure 1C). These assays revealed the presence of several complexes including those corresponding to the predicted molecular weight of NlpE or N-terminal domain dimers (labelled “D” in Figure 1C). However, we also observed the presence of several other unidentified higher molecular weight bands.

**Figure 1.**
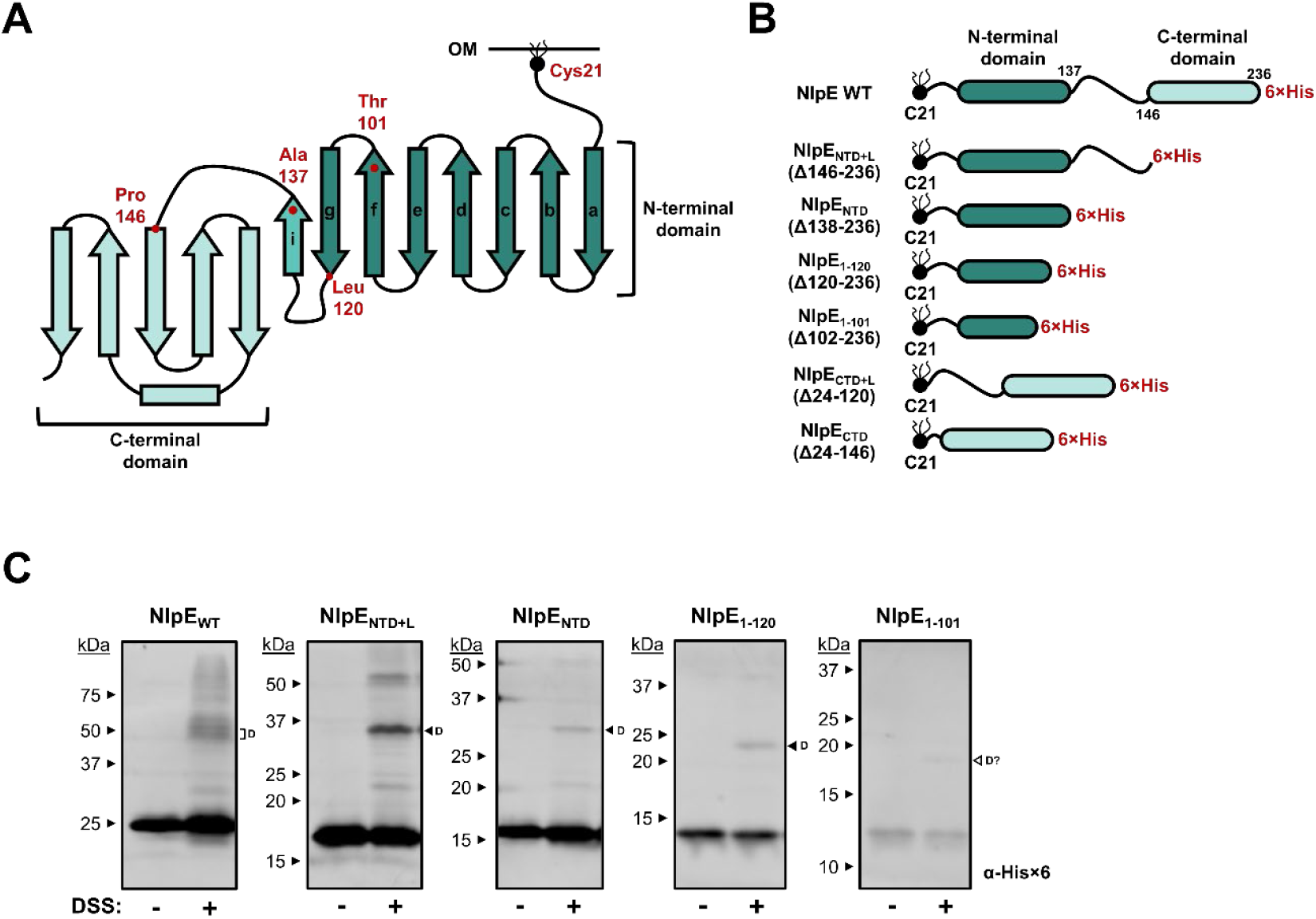
NlpE variants used in this study. **(A)** The structure of NlpE based on the hypothetical monomer model proposed by Hirano and colleagues (11). β-strands (with directionality) are represented as arrows whereas α-helices are represented without arrowheads. The β-strands making up the N-terminal domain are lettered a-i. **(B)** Representations of the NlpE variants expressed from various plasmids used in this study. The amino acid number includes the signal peptide (i.e. the lipid-modified N-terminal cysteine is labelled Cys21 not Cys1). **(C)** *In vivo* DSS crosslinking of cells expressing His-tagged NlpE variants containing the N-terminal domain. NlpE variants were expressed via leaky expression from pTrc99A and then subjected to crosslinking. Samples were analyzed by Western blotting with anti-His×6 antibody. “D” indicates putative NlpE dimers. The “D?” indicates a band present in NlpE_1-101_-expressing cells that appears to be too small to be confidently identified as a dimer.

To study these novel NlpE-containing complexes, we conducted a pull-down assay utilizing purified, soluble, C-terminal His×6-tagged NlpE to capture potential interaction partners from *E. coli* membrane preparations. The resulting complexes were separated by SDS-PAGE and proteins were visualized by silver staining (Figure 2A). Two prominent bands of around 27 and 48 kDa (red arrowheads in Figure 2A) were observed which appear to correspond to an NlpE monomer and dimer and are observed even in denaturing conditions (see also Figure S1B). A ∼30 kDa band (white arrowhead in Figure 2A) also pulled down with purified NlpE. Mass spectrometry analysis of this 30 kDa band revealed the presence of peptides corresponding to both NlpE and the major OM protein OmpA (Table S4).

**Figure 2.**
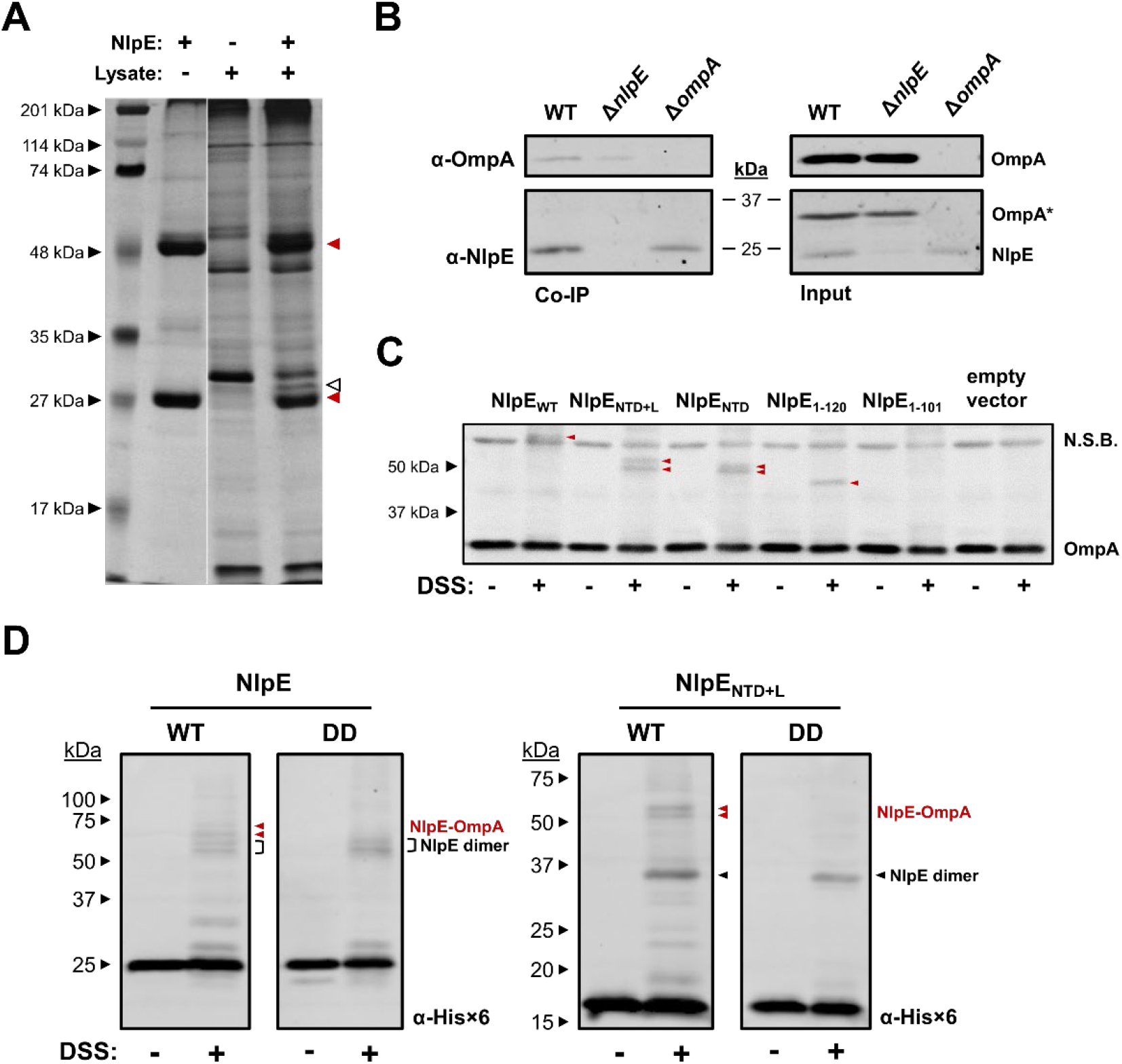
NlpE interacts with the OM protein OmpA. **(A)** Pull-down assays with His-tagged NlpE immobilized on cobalt resin and cell lysates prepared from *E. coli* MC4100. Pulled-down proteins were analyzed by SDS-PAGE and silver staining. Bands of interest were identified by mass spectrometry. Monomer and putative dimer bands of NlpE are indicated with red arrowheads, whereas pulled-down OmpA is indicated with the unfilled arrowhead. **(B)** *in vivo* co-immunoprecipitation assays confirm NlpE-OmpA interactions at native and over-produced expression levels of NlpE. Eluted proteins were analyzed by SDS-PAGE and Western blotting with anti-NlpE and OmpA antibodies and anti-PoxB was used as a specificity control. **(C)** NlpE-OmpA complexes observed by *in vivo* DSS crosslinking. Cells expressing NlpE and variants from pTrc99A were subjected to crosslinking with 0.5 mM DSS for 30 minutes. Samples were analyzed by Western blotting with anti-OmpA antibody. Resulting OmpA-NlpE complexes are indicated with red arrowheads. N.S.B. indicates a non-specific band detected by the anti-OmpA antibody. **(D)** *in vivo* crosslinking with WT and IM NlpE (DD) variants. NlpE variants were expressed from a P*ara* promoter on pBAD18 with 0.2% L-arabinose for 1 hour and cells were subjected to crosslinking with 1 mM DSS for 30 minutes. Samples were analyzed by Western blotting with anti-His×6 antibody.

To confirm NlpE-OmpA interactions *in vivo*, we conducted co-immunoprecipitation experiments on cells expressing NlpE at native levels (Figure 2B). NlpE-specific antisera was immobilized on protein A agarose beads and treated with crude membrane extracts from wildtype, Δ*nlpE*, and NlpE-overexpressing cells. We found that OmpA co-immunoprecipitated with NlpE in WT cells, despite low levels of natively expressed NlpE (Figure 2B). However, we also detected a fainter OmpA band in the co-IP fraction of Δ*nlpE* cultures. This appears to be due to cross-reaction of our anti-NlpE antisera with OmpA as confirmed by bands corresponding to OmpA in WT and Δ*nlpE* but not Δ*ompA* membrane preparations (Figure 2B, input lanes). Despite this, less OmpA was pulled down in Δ*nlpE* vs WT lysates, suggesting that NlpE can interact with OmpA at native levels of expression. Furthermore, the cross-reactivity of the anti-NlpE antisera with OmpA likely originates from OmpA which co-purified with NlpE that was used for antisera generation, providing further evidence for interactions between these two proteins.

We then performed the *in vivo* crosslinking experiments with WT NlpE and N-terminal domain containing NlpE truncation variants (Figure 1, 2C) while immunoblotting for OmpA to confirm NlpE-OmpA interactions and determine which NlpE domains contribute to this interaction. We confirmed that WT NlpE crosslinks to OmpA and observed that the N-terminal domain of NlpE is sufficient to mediate NlpE-OmpA interactions (Figure 2C). Complexes were not observed when the N-terminal domain was disrupted by the deletion of the most C-terminal beta strand (g in Figure 1A) of NlpE (NlpE_1-101_). However, this may be due to lower levels of expression of NlpE_1-101_ as this construct is significantly less stable than variants containing an intact N-terminal domain (Figure S1A); as expected, deletions extending into the folded NTD resulted in a decrease in apparent stability. To confirm these results, we repeated the experiment in an *ompA* knock-out strain, where no complexes were detected (Figure S2A). Cross-linking experiments in using cells expressing NlpE_WT_, NlpE_NTD+L_, and NlpE_NTD_ resulted in two distinct NlpE-OmpA bands (Figure 2C). These dual bands are somewhat obscured by a non-specific band when NlpE_WT_ is expressed but are clearly observable in blots with cleared anti-sera (Figure S2B).

We wondered if the NlpE C-terminal domain can interact with OmpA independently of the N-terminal domain; DSS crosslinking on strains expressing His-tagged NlpE C-terminal domain with linker (NlpE_CTD+L_) and NlpE C-terminal domain only (NlpE_CTD_) with the native NlpE signal sequence and lipobox sequence failed to produce clear evidence for NlpE_CTD_-OmpA complexes (Figure S2C). While we did observe a very faint crosslinked band when the NlpE_CTD+L_ construct was expressed, no bands were seen when expressing the C-terminal domain on its own. Thus, it’s likely that the N-terminal domain of NlpE, not the C-terminal domain, is mostly responsible for the interaction between NlpE and OmpA. Furthermore, we found that bands corresponding to NlpE-OmpA complexes are absent when expressing a permanently IM-localized variant of NlpE (Figure 2D), ruling out the possibility that IM localized NlpE interacts with OmpA. Taken together, these results place NlpE-OmpA complexes in the OM where the NlpE N-terminal domain is proximal to OmpA, rather than in a configuration where IM-localized NlpE spans across the periplasm to interact with OmpA via the C-terminal domain.

### NlpE and OmpA mediate Cpx response surface sensing

NlpE is required for activation of the Cpx response in surface-adhered cells under some conditions (6, 7). Given our discovery of NlpE-OmpA complexes, we hypothesized that these two proteins may work together to activate the Cpx response during growth on surfaces. To test this hypothesis, we conducted a surface adhesion assay where the activity of a luminescent *cpxP-lux* reporter was measured in *E. coli* MC4100 WT, Δ*nlpE*, Δ*ompA*, and Δ*nlpE*Δ*ompA* strains adhered to hydrophobic glass bead surfaces or free-floating in the media (planktonic). These assays confirmed that the Cpx response is activated in cells adhered to hydrophobic surfaces (Figure 3). The increase in Cpx activity was dependent on CpxA, confirming that the signal sensed by the system activates CpxA rather than activating CpxR directly (Figure 3A). Importantly, deletion of *nlpE* and *ompA* either individually or in combination also abolishes Cpx activation in surface-adhered cells, indicating both OmpA and NlpE are required to activate the Cpx response when cells are adhered to surfaces (Figure 3B). Taken together with our studies of NlpE-OmpA interactions, these results suggest that the NlpE-OmpA complex functions to mediate activation of the Cpx response in response to surface signals from the OM.

**Figure 3.**
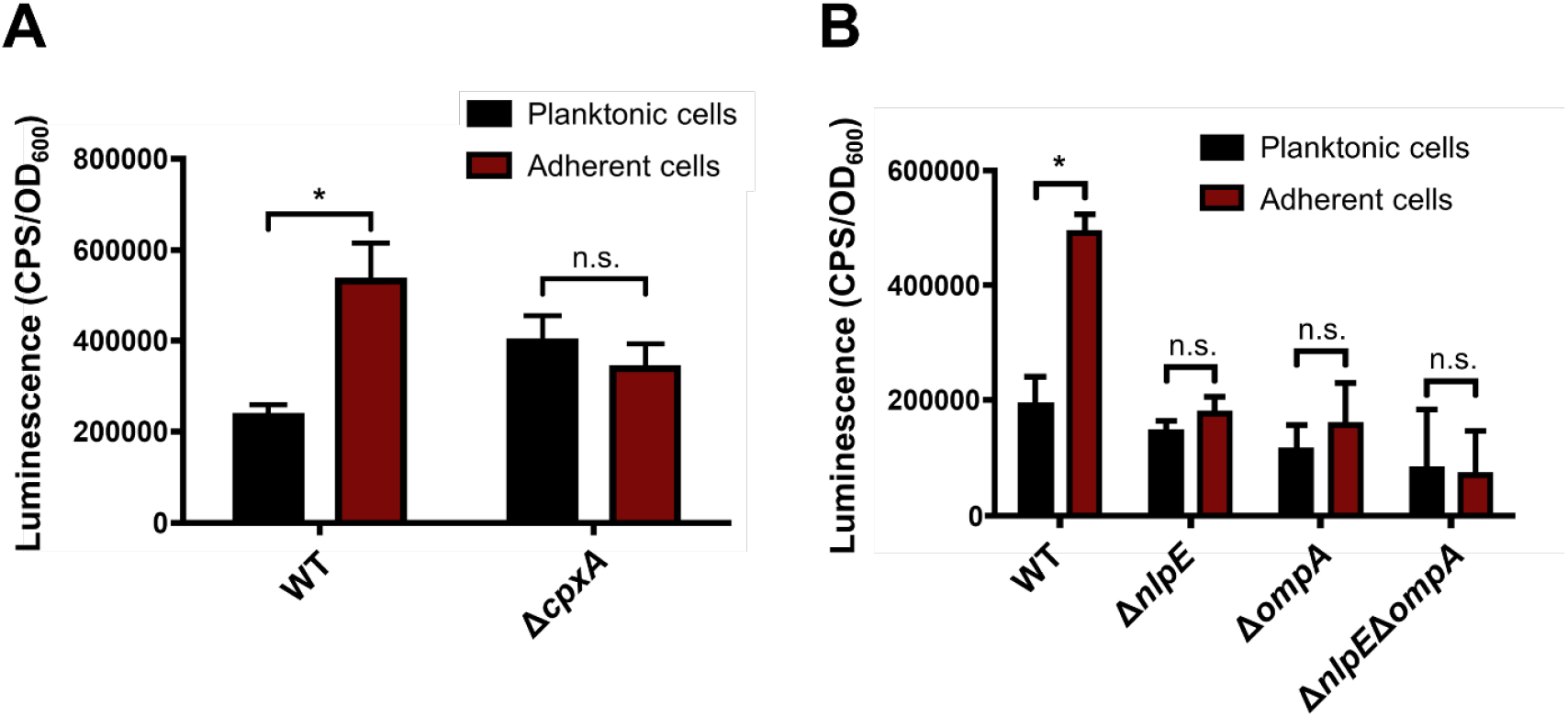
NlpE and OmpA activate the Cpx response in surface-adhered cells. Activity of the Cpx pathway upon surface adherence in **(A)** MC4100 wildtype (WT), and a Δ*cpxA* mutant and **(B)** in Δ*nlpE*, Δ*ompA*, and Δ*nlpE*Δ*ompA* strains. Overnight cultures of each strain were added to acid-washed glass beads treated to make them hydrophobic and incubated for 6h without shaking to allow adhesion. Planktonic cells were harvested from LB medium and adhered cells were collected by vortexing the beads. The activity of the Cpx pathway in planktonic cells and adhered cells was analyzed by measuring luminescence produced from a Cpx-regulated *cpxP-lux* reporter. Experiments were performed in triplicate at least three times. Shown are mean *cpxP-lux* activity values standardized to OD600 with standard deviations (one-way ANOVA test, n.s. p>0.05, *p<0.05).

### NlpE senses OmpA overexpression

OmpA protein levels are increased in both pathogenic and lab strains of *E. coli* grown in surface-adhered biofilms (12). Because both OmpA and NlpE are implicated in Cpx signaling in surface-adhered cells, we wanted to test if NlpE is sensitive to the level of OmpA in the envelope. We found that overexpression of OmpA from a plasmid activates the Cpx response in WT cells, but this activation is strongly reduced in the absence of NlpE (Figure 4). We wondered if NlpE senses OMP overexpression in general or if the signal sensed is specific to NlpE-OmpA interactions. To test this, we repeated these experiments with two sets of OMPs: larger OMPs that fold into 16-stranded β-barrels (OmpC, OmpF, and LamB; Figure 4A) and smaller OMPs that fold into 8-10 stranded β-barrels (OmpX, OmpW, and OmpT; Figure 4B). While overexpression of these OMPs lead to activation of the Cpx response to differing extents, Cpx activation in all cases, except for overexpression of OmpA and OmpX occurred independently of NlpE. Therefore, the signal sensed by NlpE is unlikely to be a general signal or stress associated with OMP overexpression.

**Figure 4.**
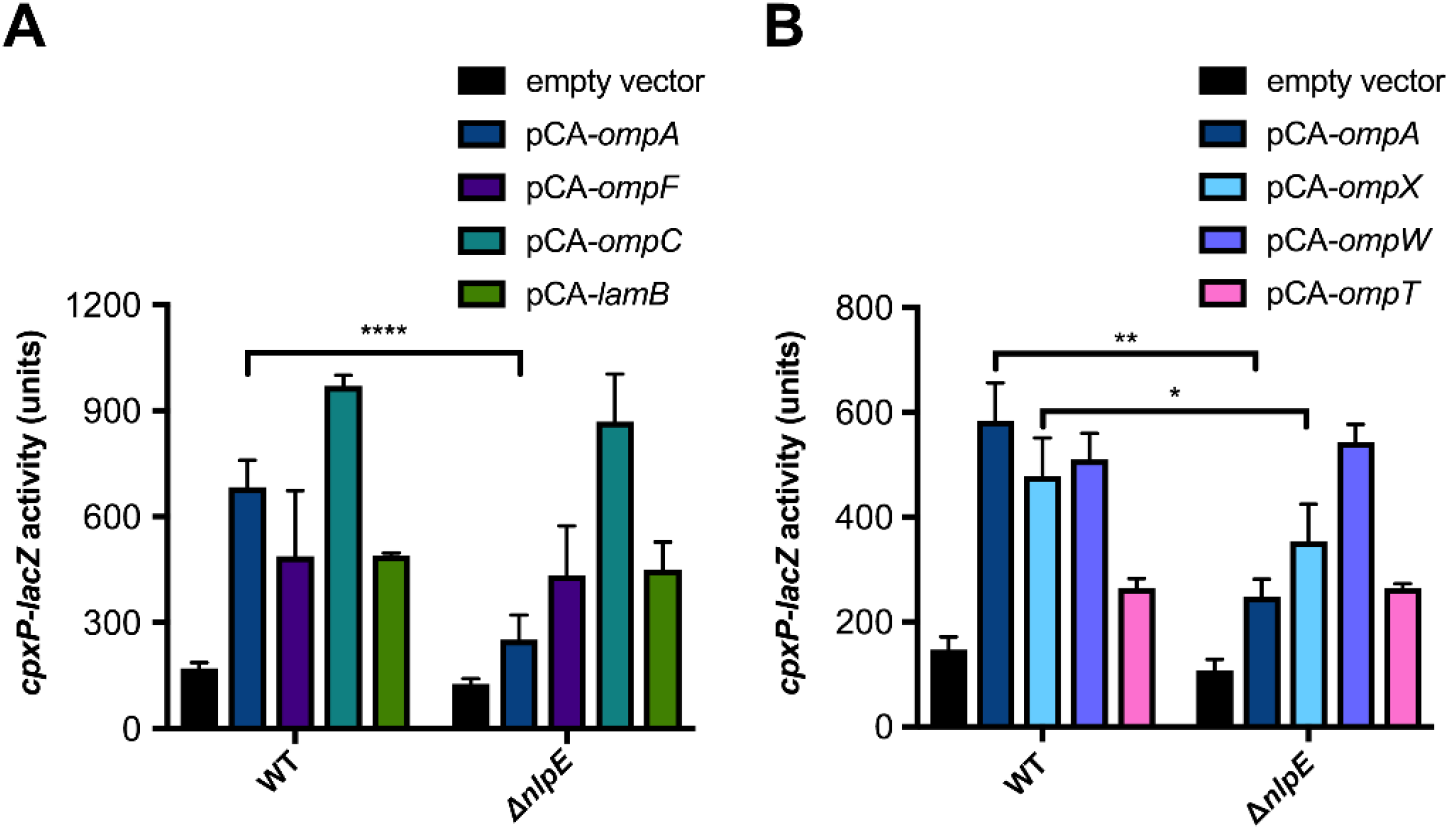
NlpE senses overexpression of OmpA but not other OMPs. OMP expression was induced with 0.1 mM IPTG in WT and Δ*nlpE* cells when cultures reached mid-log phase. Cpx activation was determined by measuring *cpxP-lacZ* activity in β-galactosidase assays. The OMPs expressed were **(A)** larger, 16-stranded and **(B)** smaller, 8-10 stranded OMPs. Shown are means with standard deviation (*t-*test, *p<0.05, **p<0.005, ****p<0.0005).

OmpA is a unique OM protein in both its structure and function. Structurally, OmpA consists of two distinct domains: an N-terminal, OM-integral β-barrel domain and a C-terminal globular periplasmic domain with a proline-rich unstructured linker that connects these two elements (13, 14). OmpA is also thought to adopt a larger pore conformation, although the physiological circumstances under which this form occurs remain unclear (15, 16). OmpA also appears to dimerize via C-terminal domain to C-terminal domain interactions (17, 18). In order to further investigate how NlpE senses OmpA overexpression, we created vectors expressing variants of OmpA by site-directed mutagenesis. These variants lack the C-terminal domain all together (OmpA_ΔCTD_) or contain an alanine substitution for the K213 lysine residue essential for dimerization (17, 18). While the OmpA K213A variant is expressed comparably to WT OmpA, levels of the OmpA_ΔCTD_ construct were lower than wildtype, suggesting this variant is unstable (Figure S3). Despite this, we observed that the Cpx response is activated to similar levels in both these backgrounds, albeit at lower overall levels compared to when the wildtype OmpA is overexpressed (Figure 5A). However, unlike when overexpressing WT OmpA, a significant decrease in Cpx activity was not seen in the absence of NlpE suggesting that both the OmpA C-terminal domain and OmpA dimerization are necessary for NlpE signaling from the OM.

**Figure 5.**
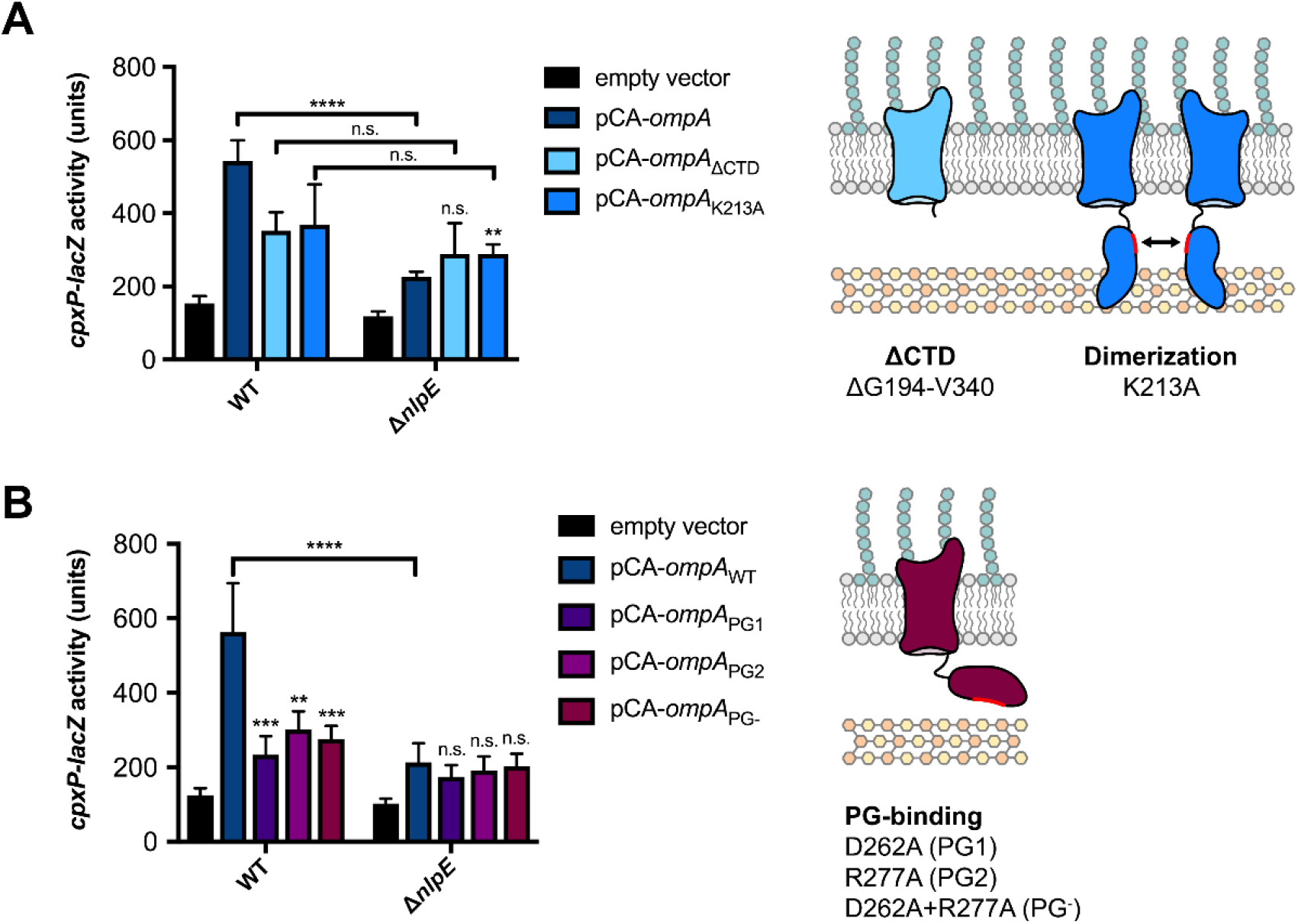
The OmpA C-terminal domain influences Cpx activation through NlpE. The ability of WT OmpA and variants that **(A)** lack the periplasmic C-terminal domain or are deficient in dimerization (K213A) or **(B)** are mutated in residues involved in peptidoglycan binding to activate the Cpx response in WT and Δ*nlpE* cells. OmpA expression was induced with 0.1 mM IPTG for 1 hour after cultures reached mid-log phase of growth. Shown are mean *cpxP-lacZ* activities with standard deviations (*t-*test, **p<0.01, ***p<0.005, ****p<0.0001). Significance markers shown directly above bars indicate tests comparing reporter activity in that strain to that of same strain background expressing WT OmpA.

The OmpA C-terminal domain non-covalently binds the peptidoglycan (PG) cell wall, thus regulating the spatial characteristics of the envelope and providing structural support (19, 20). Two charged residues located in the C-terminal domain are important for PG binding: D262 and R277. These residues were determined in the OmpA of *Acinetobacteri baumanii* (19); however, these residues are highly conserved across Gram-negative bacteria, including *E. coli*. We mutated these two residues individually (named PG1 and PG2) and in combination (PG-) and determined the ability of these variants to activate the Cpx response through NlpE when overexpressed. We observed that mutating any of the OmpA PG-binding residues to alanine is sufficient to significantly reduce activation of the Cpx response upon overexpression in WT cells to a similar extent as deleting *nlpE* (Figure 5B), despite these mutants being expressed at similar levels to WT (Figure S3). Expressing these variants in a Δ*nlpE* strain resulted in comparable levels of activation to WT (1.9-2.4 fold in WT vs. 1.7-2.0 fold in Δ*nlpE*) suggesting that these mutations impact ability of NlpE to sense the OmpA variant being expressed. Furthermore, we did not observe significantly different levels of Cpx induction when expressing WT OmpA or the PG binding mutants in the Δ*nlpE* background. Thus, the OmpA C-terminal domain, and in particular its ability to bind the cell wall and mediate dimerization, appears to play a key role in activating the Cpx response through NlpE when OmpA is overexpressed.

### The C-terminal domain of NlpE mediates Cpx activation from the OM

Because the N-terminal domain of NlpE interacts with OmpA, we hypothesized that the C-terminal domain of NlpE mediates activation of the Cpx response from the OM. To test this, we deleted the C-terminal domain of the chromosomal *nlpE* locus by allelic exchange and tested if this chromosomally expressed NlpE_ΔCTD_ variant (equivalent to NlpE_NTD+L_) can sense OmpA overexpression. We found that NlpE_ΔCTD_ is no longer able to sense OmpA overexpression; Δ*nlpE* and *nlpE*_ΔCTD_ strains showed significantly reduced Cpx activation during OmpA overexpression, and overexpressing OmpA lead to similar levels of fold-activation in Δ*nlpE* (2.4 fold) and *nlpE*_ΔCTD_ (1.9 fold) strains as opposed to WT (4.0 fold) (Figure 6A). Thus, while the N-terminal domain interacts with OmpA, the C-terminal domain appears to be responsible for transmitting this signal to CpxA. Our data from overexpressed NlpE variants suggests that NlpE lacking the C-terminal domain is stably expressed (Figure S1A); natively expressed NlpE_ΔCTD_ is also expressed comparably to WT (Figure S4), suggesting that our results are not due to differences in NlpE expression level.

**Figure 6.**
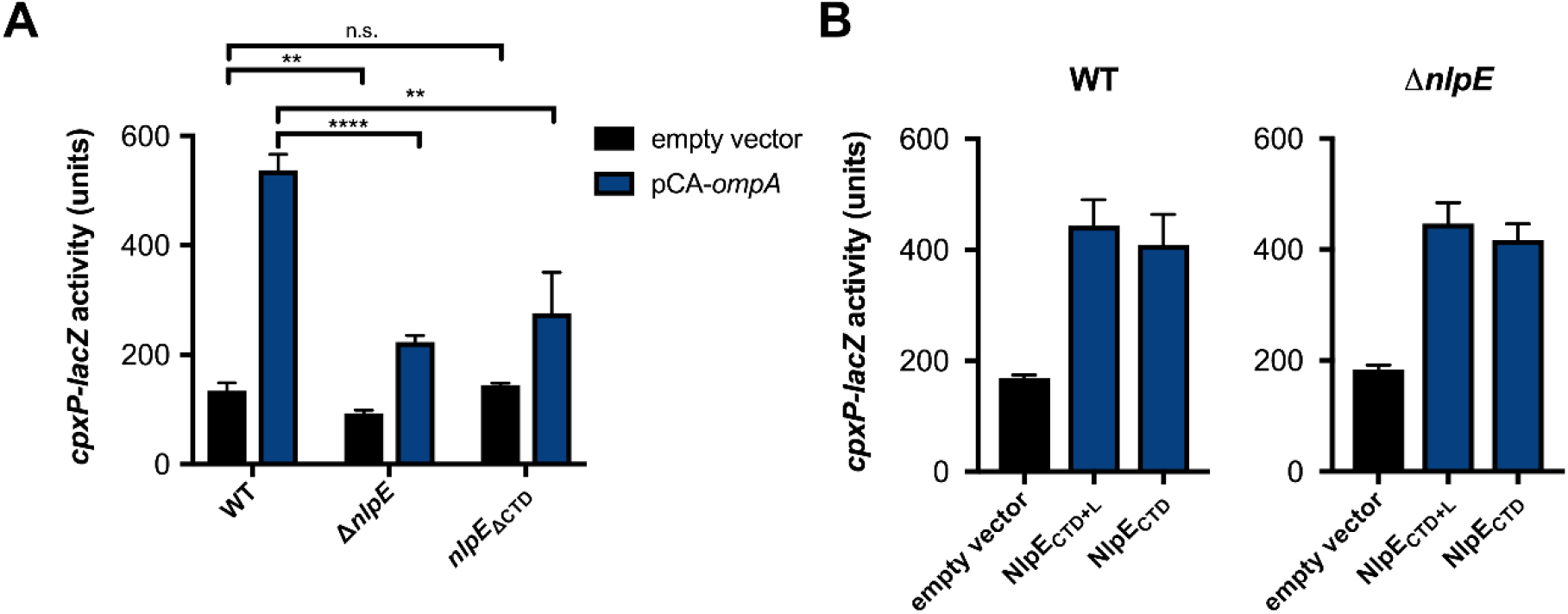
The C-terminal domain mediates NlpE signaling from the OM. **(A)** The ability of OmpA overexpression to activate the Cpx response in WT, Δ*nlpE*, or nlpE_ΔCTD_ (NlpE with the C-terminal domain deleted). **(B)** The ability of NlpE lacking the N-terminal domain (NlpE_CTD+L_) or lacking the N-terminal domain and linker (NlpE_CTD_) to activate the Cpx response when overexpressed in WT and Δ*nlpE* TR50. For (A) and (B), mean *cpxP-lacZ* activities with standard deviations are shown (*t-*test **p<0.01, ****p<0.0001).

We also found that overexpressing variants of NlpE that lack the N-terminal domain leads to activation of the Cpx response, albeit to a much weaker extent than overexpressing the N-terminal domain (Figure 6B). This weaker activation, however, may be due to comparatively lower levels of expression of these constructs (Figure S1A). To rule out the possibility that overexpressing these constructs decreases the efficiency of lipoprotein biogenesis, causing WT NlpE to activate CpxA at the IM, we repeated these experiments in an Δ*nlpE* mutant and found the same results. Overall, these results suggest that the C-terminal domain can activate the response independently of the N-terminal domain.

## Discussion

NlpE senses OM lipoprotein trafficking defects (8-10) and has also been demonstrated to activate the Cpx system in response to surface adhesion (6, 7). In this study, we reveal that NlpE does not function alone in this latter signaling role, but cooperates with the major OMP OmpA and utilizes its distinct two-domain structure to mediate activation of the Cpx ESR connected to OM-associated signals (Figure 7).

**Figure 7.**
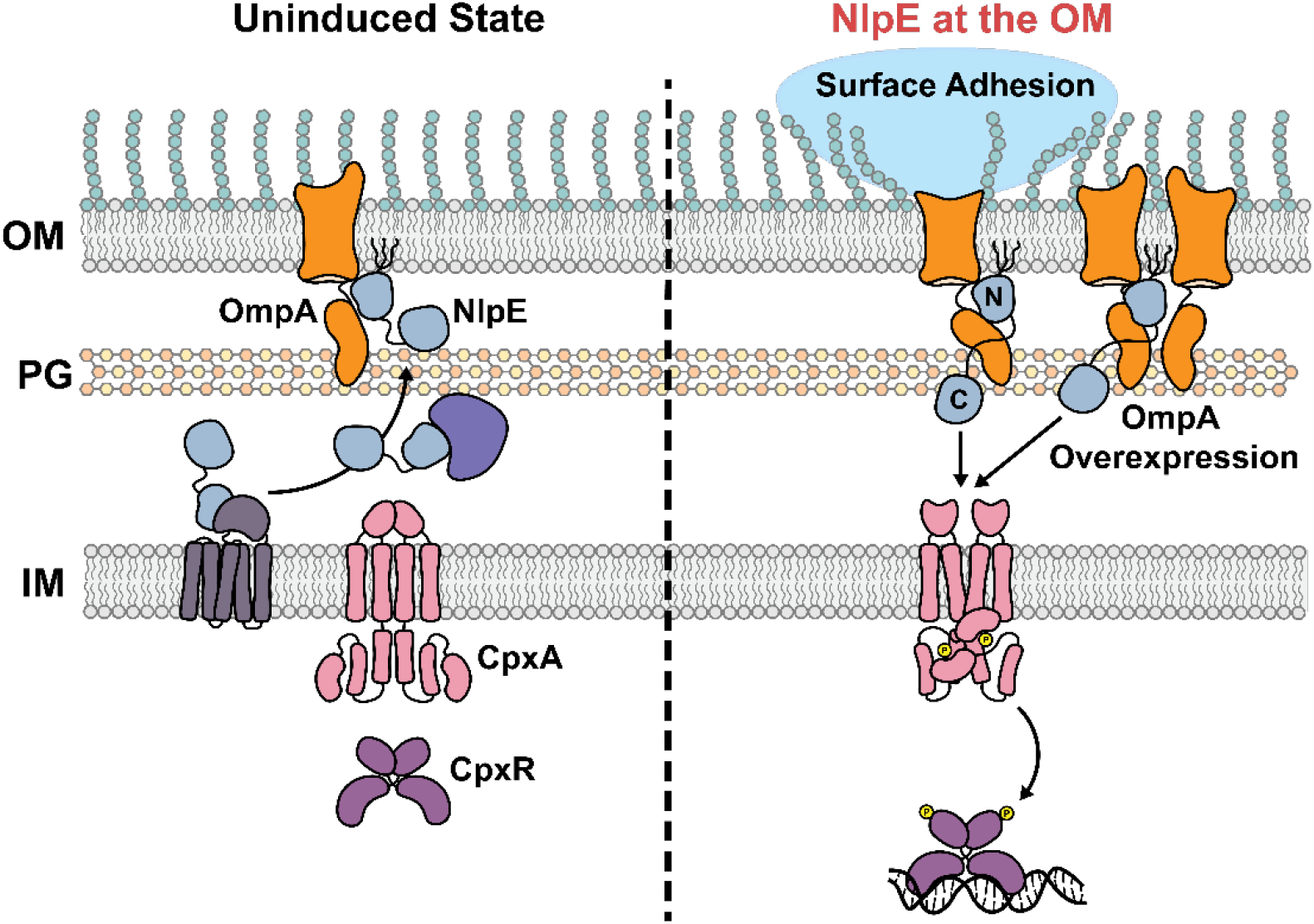
NlpE is an OmpA associated OM sensor. In the absence of inducing cues, NlpE is processed and trafficked efficiently to the OM where it forms complexes with OmpA via the NlpE N-terminal domain. In the presence of a surface-related cue, such as surface adhesion, or OmpA overexpression, NlpE and OmpA cooperate to transduce a signal to CpxA via the NlpE C-terminal domain (left panel). Whether this activation occurs by direct interaction or is mediated by other factors is currently unknown.

### NlpE is an OmpA-associated sensor

The mechanisms of NlpE signaling from the OM have remained largely mysterious. Several studies have implicated OmpA in activating the Cpx response under different conditions, but no clear mechanisms have been proposed (21, 22). Our study reports that NlpE interacts with OmpA to signal certain OM signals to the Cpx response. That NlpE signals from the OM and not the IM is made clear by the observations that the N-terminal domain, not the C-terminal domain, of NlpE interacts with OmpA (Figures 2C, S2C) and that permanently IM-bound NlpE does not crosslink to OmpA (Figure 2D). These findings imply that direct N-terminal domain-CpxA interactions do not mediate activation of the response in this context. The fact that deleting the C-terminal domain greatly diminishes induction of the Cpx response during OmpA overexpression provides support for this interpretation and strong evidence for a model where signals sensed by NlpE in the OM in conjunction with OmpA are transduced through the C-terminal domain to induce the Cpx response.

Our findings naturally draw comparisons to RcsF, another lipoprotein activator of an envelope stress response (i.e. the Rcs phosphorelay) that interacts with OmpA (23-30). While RcsF can be folded into the OmpA β-barrel, as evidenced by the requirement for unfolded OmpA for complex formation *in vitro* (23), our pull-down assays suggest that NlpE interacts with folded OmpA (Figure 2A). Dekoninck and colleagues report that the globular OmpA C-terminal domain interacts directly with RcsF’s C-terminal domain and propose a buffer mechanism where OmpA competes with IgaA for RcsF binding (28), although evidence contrary to this model indicates that more work is required to elucidate the precise mechanisms of OmpA-RcsF signaling (23, 29). In contrast, the NlpE-OmpA signaling interaction appears to be more cooperative than competitive in nature. While both CpxA and OmpA bind the NlpE N-terminal domain, the spatial and structural characteristics of this domain make it unlikely that CpxA and OmpA compete for binding. Furthermore, deleting OmpA does not lead to constitutive activation of the response (Figure 3B) as would be expected if release of NlpE from OmpA is responsible for Cpx activation. However, it is possible that this is because NlpE may interact with other OMPs; more work is required to determine if NlpE, like RcsF, associates with multiple OMPs.

Instead, the requirement of the NlpE C-terminal domain for signaling OmpA overexpression supports a model where signals are transduced through this domain to activate the Cpx response, akin to the models of signaling proposed in the original structural study (11). However, the precise mechanisms by which the C-terminal domain accomplishes this remain unclear. Hirano and colleagues propose that partial unfolding of the N-terminal domain allows for the linker region to extend into the periplasm enough for the C-terminal domain to directly interact with CpxA (11). Previous studies suggest that the NlpE C-terminal domain is not needed for direct interaction with the CpxA sensor domain (10), and we have also failed to observe evidence for NlpE C-terminal domain-CpxA interactions in two-hybrid assays (in preparation), thus suggesting that another mechanism besides direct-physical interaction may mediate CpxA activation by this domain. Although not examined in this study, it is possible that the redox state of the disulfide bond present on the C-terminal domain may be involved in activation of the Cpx response in this situation as suggested in a previous study (10). Alternatively (or perhaps in conjunction), other envelope proteins may mediate NlpE signaling from the OM. We observed that several crosslinked products are only present when overexpressing OM-localized NlpE compared to permanently IM-bound NlpE (Figure 4D). A far more expansive network of envelope proteins than what is currently known may be involved in mediating signal transduction across the envelope.

We observed that overexpression of OmpA lacking the C-terminal domain results in low level activation (less than two-fold) of the Cpx response independently of NlpE (Figure 5. The C-terminal globular domain of OmpA is involved in a number of different functions including homodimerization (17) and cell wall binding (19). Interestingly, we found that, similarly to deleting the C-terminal domain, mutating amino acids associated with these functions leads to lower levels of Cpx activation and abolishes the dependence of the remaining activation on NlpE (Figure 5). There is controversy around the exact conformation of OmpA *in vivo;* some studies suggest that switching between small pore (presumably with a globular C-terminal domain) and large pore conformations is possible (15, 16, 31). Furthermore, portions of the OmpA C-terminal domain can become surface exposed in *Salmonella enterica* (32). However, the fact that altering functions associated with the globular C-terminal domain impacts signaling through NlpE suggests that NlpE may interact with the small-pore, periplasmic domain-containing conformation of OmpA rather than the larger pore form. As the precise region of OmpA required for NlpE interaction were not determined in this study, further work is required to further characterize NlpE-OmpA interactions.

Molecular dynamics simulation studies of OmpA suggest that OmpA dimerization may allow the C-terminal domain to bind the cell wall at a distance from the OM (33). Thus, abolishing OmpA dimerization may alter the properties of OmpA binding to peptidoglycan. Mutating the two conserved PG-binding residues in the C-terminal domain of OmpA, either individually or in combination, significantly reduces activation of the Cpx response during overexpression (Figure 5B). The PG-binding property of OmpA may, therefore, have an important role in NlpE signaling from the OM. The spatial characteristics of the periplasm, namely the distance between the OM and IM, is a crucial factor in allowing RcsF to signal across the envelope (34). However, one of the mysteries of signaling across the periplasm is how proteins in the OM or the IM “cross” the cell wall to communicate with proteins in other compartments. The association between signaling proteins such as NlpE and PG-binding proteins such as OmpA may function to overcome this challenge by allowing OmpA to “guide” OM-localized signaling proteins across the cell wall where they can access the IM. Further investigation of PG-binding envelope proteins and their role in signal transduction may shed light on how signals are communicated across physically segregated spaces in the envelope.

### NlpE and OmpA mediate adaptation to surfaces

Bacterial surface adhesion and biofilm growth are complex processes influenced by many factors including nutrient availability, mechanical stimuli, stress responses, and small molecule signaling (reviewed in (35, 36)). Some evidence has implicated the Cpx response in sensing surfaces (6, 7, 21) and regulating processes related to surface adhesion and growth (21, 37-44). However, the precise role of the Cpx response in mediating adhesion and surface growth appears to be quite complex. The Cpx response regulates the expression and assembly of adhesion-related appendages such as curli (37), and chaperone-usher and type IV pili (45-48) while also downregulating motility (7, 49, 50). Overexpression of NlpE increases biofilm production through the Cpx-regulated diguanylate cyclase DgcZ (44), and deletion of *cpxR* and *nlpE* diminishes attachment to hydrophobic surfaces (6).

Our study suggests that NlpE-OmpA complexes are a part of the mechanism by which the Cpx response responds to signals related to surface growth. OmpA and its homologues are prominent Gram-negative adhesins and invasins involved in the pathogenesis of several Gram-negative species (reviewed in (51)). In agreement with emerging literature, our results expand our understanding of OmpA not only as a virulence factor with direct implications for host cell adhesion and invasion, but also as a signaling factor mediating the transduction of surface-related signals to regulatory systems such as the Cpx and Rcs envelope stress responses. Deleting *ompA* is sufficient to abolish activation during surface adhesion, suggesting that NlpE interacts primarily with OmpA to mediate signaling unlike RcsF, which appears to interact with several OMPs in a similar capacity (23). While defects in OMP biogenesis are a classical inducing cue of the σ^E^ response, we find that overexpression of several OMPs induces the Cpx response. This is in line with emerging studies supporting the conclusion that the Cpx response may respond to defective OMP biogenesis as a means to protect the IM; the small regulatory RNA (sRNA) CpxQ downregulates the expression of the chaperone Skp, preventing BAM-independent incorporation of OMPs into the IM (52, 53). Similarly, we speculate that the induction of the Cpx response by OMP overexpression observed in this study (Figure 4), may increase Skp-dependent IM incorporation, thereby activating the Cpx response. In support of this, we noticed that overexpression of OMPs such as OmpA, OmpC, OmpF, and LamB lead to similar growth defects (Figure S5), suggesting aberrant expression of these proteins generates significant stress for cells. Overall, this conclusion aligns with the finding that Cpx ESR activation by most OMPs is not NlpE dependent, and that OmpA overexpression residually induces the Cpx pathway in absence of NlpE (Figure 4).

The uniqueness of the NlpE-OmpA interaction is underscored by the observation that NlpE does not sense the overexpression of other OMPs in a similar manner to OmpA (Figure 4). Furthermore, if OMP overexpression does lead to increased IM stress, the fact that this stress is sensed independently of NlpE further reinforces its role as an OM-specific sensor. A previous study suggested that OmpA production leads to increased envelope stress, activation of the Cpx response, and changes to biofilm formation via regulation of cellulose production (21). Our results suggest that, NlpE cooperates with OmpA to activate the Cpx response in surface-adhered cells. While OMP overexpression appears to lead to envelope stress as indicated by an extended lag phase in growth curves (Figure S5), it is unlikely that NlpE senses all of the signals generated by OmpA overexpression as deleting *nlpE* does not significantly alter growth and overexpressing other OMPs leads to similar growth phenotypes. Thus, NlpE and OmpA operate together in a signal transduction pathway to mediate Cpx activation in the presence of a surface-specific related inducing signal.

The precise nature of this surface signal remains mysterious as it is unclear whether NlpE senses surfaces themselves or some cue present in surface-adhered/biofilm cells. A recent study has questioned the role of the Cpx response in direct surface sensing based on the finding that Cpx-regulated fluorescent reporters are not activated in cells grown in flow chamber surfaces (54). Thus, it is possible that the Cpx response does not directly sense surfaces themselves, but is activated in response to some other signal present in surface-adhered cells. It has previously been reported that OmpA levels are increased, both in clinical and nonclinical strains of *E. coli*, when cells are grown in surface-adhered biofilms (12). We report that NlpE is sensitive to OmpA overexpression, suggesting that the level of OmpA in the OM may be part of the mechanism by which OmpA and NlpE mediate activation of the Cpx response in adhered cells (12). Nevertheless, the Cpx response is differentially induced by the type of the surface (e.g. hydrophobic vs. hydrophilic) (6, 7), and OmpA itself is a major surface-exposed antigen of Gram-negative bacteria with roles in adhesion and biofilm formation (51), both suggesting a direct surface-sensing role for the Cpx response through complexes of NlpE and OmpA.

## Conclusions

Recent studies of RcsF clearly establish close links between OM lipoprotein and OMP biogenesis with significant consequences for bacterial function (23-30). Our findings show that RcsF is not exceptional in this regard. The implication of OmpA in Cpx response signaling through NlpE further expands our understanding of OmpA as a novel signal transduction factor that regulates key adaptive systems and the complex interdependence of envelope biogenesis pathways that facilitate stress adaptation. Further study of these envelope signaling complexes will be key to understanding how bacteria respond to stress as well as sense and adapt to the environmental cues essential for *in vivo* success.

## Materials and Methods

### Bacterial strains, growth, and strain construction

The strains used in this study are listed in Table S1. All strains were grown in lysogeny broth (LB) at 37°C with shaking at 225 RPM that was supplemented with the following concentrations of antibiotics as appropriate: ampicillin (100 µg/ml), kanamycin (50 µg/ml), chloramphenicol (25 µg/ml), tetracycline (12 µg/ml), and spectinomycin (25 µg/ml). To induce expression from inducible promoters, 0.1 mM isopropyl β-d-1-thiogalactopyranoside (IPTG) or 0.2% L-arabinose was added to cultures that were grown to an optical density of 0.4-0.6 (A_600_) for 0.5-2 hours depending on the experiment.

All whole gene deletion mutants were constructed by P1 transduction using lysates derived from the corresponding mutants from the Keio library (55). Kan^R^ cassettes were removed using FLP-mediated recombination as previously described (56). *nlpE* chromosomal mutants were created by allelic exchange. To create TR50 *nlpE*_ΔCTD_, 1 kb up and downstream of *nlpE* was amplified by PCR, and Gibson assembly (New England Biolabs) was used to recombine these fragments into pRE112 digested with XbaI and PaeI. Gibson reaction products were transformed into OneShot PIR1 competent cells (Thermo Fisher). Allelic exchange vectors(57) were sequenced prior to conjugation (Molecular Biology Facility, University of Alberta). Suicide vectors were then transformed into the donor strain MFDλpir and mated with the recipient TR50 strains on LB plates with 0.3 mM diaminopimelic acid (DAP). Transconjugants were selected by plating on LB with chloramphenicol and incubating overnight. To select for double crossovers, a chloramphenicol-resistant colony was grown in LB with no antibiotics for 6 hours and plated onto LB without NaCl with 5% sucrose and incubated at room temperature for two nights. Sucrose-resistant colonies were screened by PCR to identify colonies with the correct mutation.

### Expression vector construction and site-directed mutagenesis

The plasmids used in this study are listed in Table S2. Overexpression plasmids were created by restriction digest cloning utilizing standard procedures and the primers listed in Table S3. Site-directed mutagenesis was used to introduce deletions and substitutions into expression vectors using the Q5 site directed mutagenesis kit (New England Biolabs) and according to manufacturer’s instructions. The primers used for site-directed mutagenesis are listed in Table S3. For consistency, the numbering of amino acids in envelope proteins throughout this study includes the amino acids of the signal peptide. The sequence of inserts/mutants were confirmed by sequencing (Molecular Biology Facility, University of Alberta).

### β-galactosidase assays

*cpxP-lacZ* activity and cAMP production during two-hybrid assays were measured by quantifying β-galactosidase activity as previously describe (3, 58). Briefly, *E. coli* TR50 (MC4100 *cpxP-lacZ*) were grown overnight in LB and then subcultured in 2 ml LB. Cultures were spun down and cell pellets were resuspended in Z-buffer (59). The optical density (OD at A_600_) was measured and then each culture was treated with chloroform and sodium dodecyl sulfate (SDS) and vortexed to release β-galactosidase. β-galactosidase activity was quantified by measuring the A_420_ of each culture 25 times at 30s intervals after the addition of 10 mg/ml ortho-nitrophenyl β-galactoside (ONPG). *cpxP-lacZ* activity was calculated as the maximum slope of the linear region of A_420_ measurements standardized to that culture’s OD. The statistical significance was calculated using unpaired *t-*tests (Prism, Graphpad).

### *In vivo* DSS crosslinking

*In vivo* crosslinking with the membrane permeable crosslinker disuccinimidyl suberate (DSS; Thermo Scientific) was conducted as previously described with minor modifications (60). The optical density (A_600_) of cultures was used to collect a standardized amount of cells corresponding to an OD 2.0 in 200 μl. Cells were washed four times with phosphate buffered saline (PBS) and then subjected to crosslinking with 0.5 or 1 mM DSS for 30 minutes at room temperature. 5 μl of 1 M Tris-HCl was added to quench any excess crosslinking reagent for 5-10 minutes. Cells were then washed one more time with PBS. Cell lysates were prepared by resuspending pellets in MilliQ H_2_O and 2×Laemmli sample buffer (Sigma) and heating at 95°C for 5 minutes. SDS-PAGE and Western blotting was used to visualize crosslinked complex formation.

### Luminescence reporter assay in surface adhered cells

Activation of the Cpx envelope stress response upon adherence of bacteria to hydrophobic glass beads was measured using a Cpx-regulated *cpxP-lux* luminescent reporter gene, as previously described (61), and modifications of an assay developed by Otto & Silhavy (6). Acid-washed glass beads (212-300 μm; cat No. G1277. Sigma) served as the abiotic surface for cells to adhere to. To make the glass beads hydrophilic, glass beads were treated in a flask with a mixture of H_2_O:H_2_O_2_:NH_4_OH (Ratio 6:1:1, V:V) at 80^°^C for 10 min followed by thorough rinsing with distilled water. Glass beads were then treated with a mixture of H_2_O:H_2_O_2_:HCl (Ratio 5:1:1, V:V) at 80^°^C for 10 min followed by thorough rinsing with distilled water. Hydrophilic glass beads were immersed in distilled water for storage at room temperature overnight. To prepare hydrophobic glass beads, hydrophilic beads were first thoroughly washed with two-volumes of 99% ethanol and trichloroethylene (TCE) and treated with a 10% dimethyldichlorosilane (DDS) solution dissolved in trichloroethylene for 10 min. Excess DDS was thoroughly washed away by sequential rinses with ethanol, TCE and ethanol. Hydrophobic glass beads were immersed in ethanol for storage at room temperature overnight.

To measure Cpx activation in surface adhered cells, 1 mL of cells cultured overnight in LB for 14-16 h were added to 3 g glass beads in a flask and left at 37^°^C for 6 h, allowing adherence to occur. Planktonic (i.e. unadhered) cells were aspirated by removing LB culture from the glass beads. Glass beads were washed three times with 1 mL of fresh LB medium. 1 mL LB was added to the glass beads. Adherent cells were collected by vortexing the glass beads for 15 sec and were transferred to a new microcentrifuge tube. The activity of the Cpx response was quantified by measuring the light produced (in counts per second, CPS) from the plasmid-based *cpxP-lux* reporter construct in both planktonic and adherent cells. This value was standardized to the optical density of the solution containing planktonic or adherent cells. Statistical significance of results was determined using a one-way ANOVA test in Prism 7 (GraphPad).

### Pull-down assays and mass spectrometry analysis

NlpE-interacting proteins were identified by mass spectrometry analysis of bands excised from silver stained gels after electrophoresis of elution fractions derived from a cobalt resin column on which NlpE was immobilized prior to incubation with membrane extracts. Purified soluble C-terminally His-tagged cytoplasmic NlpE with no N-terminal lipid modification signal bound to Cobalt resin (Pierce) was washed with wash solution (50mM sodium phosphate, 300mM sodium chloride, 1% Triton X-100; pH 7.4). Membrane preparations of *E. coli* MC4100 were prepared as previously described (62). This membrane extract was added to NlpE-bound resin and agitated overnight at 4°C. The resin was then collected by centrifugation and washed with wash solution. Pulled-down proteins were eluted using elution buffer (50mM sodium phosphate, 300mM sodium chloride, 1% Triton X-100, 150mM imidazole; pH 7.4) and separated by SDS-PAGE gel. Silver staining was performed as previously described (63).

Bands indicative of proteins pulled-down specifically with NlpE were identified by mass spectrometry (Alberta Proteomics and Mass Spectrometry Facility). Briefly, bands excised from gels were destained, reduced, alkylated, and digested with trypsin in-gel. Tryptic peptides were resolved by nanoflow high performance liquid chromatography (HPLC; Easy-nLC II, Thermo Scientific) coupled to an LTQ XL-Orbitrap hybrid mass spectrometer (Thermo Scientific). Peptide mixtures were injected onto the column at a flow rate of 3000 nL/min and resolved at 500 nL/min using 45 min linear gradients from 0 to 45% v/v aqueous ACN in 0.2% v/v formic acid. The mass spectrometer was operated in data-dependent acquisition mode, recording high-accuracy and high-resolution survey Orbitrap spectra using external mass calibration, with a resolution of 30 000 and m/z range of 400–2000. The fourteen most intense multiply charged ions were sequentially fragmented by using collision induced dissociation, and spectra of their fragments were recorded in the linear ion trap; after two fragmentations, all precursors selected for dissociation were dynamically excluded for 60 s. Data was processed using Proteome Discoverer 1.4 (Thermo Scientific) and a non-reviewed Uniprot (uniprot.org) *E. coli* database was searched using SEQUEST (Thermo Scientific). Search parameters included a precursor mass tolerance of 10ppm and a fragment mass tolerance of 0.8 Da. Peptides were searched with carbamidomethyl cysteine as a static modification and oxidized methionine and deamidated glutamine and asparagine as dynamic modifications.

### *In vivo* co-immunoprecipitation assays

Large volume cultures were growth to OD 1.0 at 37°C with shaking at 225 RPM. Cells were harvested and washed once with PBS. The optical density was standardized across all cultures at this stage (∼OD 12 in 20 ml of PBS). Cells were gently lysed using the BugBuster Master Mix reagent (EMD Millipore) with 1 tablet per sample of cOmplete Protease Inhibitor Cocktail (Roche) for approximately 2 hours. Debris and unlysed cells were pelleted by centrifugation at 15,000×g for 20 minutes. Lysates were then centrifuged at 43,000 RPM at 4°C for 45 minutes to pellet membranes. Membrane pellets were solubilized in membrane solubilization buffer (25 mM Tris-HCl pH 7.4, 150 mM NaCl, 0.5% Triton X-100) overnight with gentle rotation. Co-immunoprecipitations were conducted with the Pierce Crosslink IP Kit essentially according to manufacturer’s instructions. BCA assays (Pierce) were used to calculate sample protein concentrations; 1 µg of membrane preparations was used in assays. Co-immunoprecipitations were conducted overnight at 4°C with gentle rotation. Eluates from co-immunoprecipitations were analyzed by SDS-PAGE and Western blotting.

### SDS-polyacrylamide gel electrophoresis and Western blotting

SDS-PAGE and Western blotting was conducted according to standard protocols. Where indicated, protein concentrations of solutions were determined by BCA protein assay (Pierce) following manufacturer’s instructions. Briefly, samples were separated on 8-12% SDS-PAGE gels and transferred to nitrocellulose membranes using semi-dry transfer (BioRad Trans-Blot Semi-Dry Transfer Cell). Membranes were blocked in 5% non-fat milk in Tris-buffered saline with 0.1% Tween-20 (TBST) and probed with primary antibody overnight at 4°C in 2% BSA in TBST at the following concentrations: anti-NlpE (rabbit polyclonal, this study, 1:8,000-40,000), anti-His×6 (mouse monoclonal, Invitrogen, 1:10,000-20,000), anti-OmpA (rabbit polyclonal, Antibody Research Corporation, 1:5,000-10,000), anti-CpxA-MBP (rabbit polyclonal, (57), 1:10,000) and anti-RNAP (mouse monoclonal, BioLegend, 1:5,000). Chemiluminescent alkaline phosphatase (AP)-conjugated (BioRad) or fluorescent IRDye 800CW (goat anti-rabbit) and 680RD (goat anti-mouse) antibodies were used to detect proteins. Chemiluminescent signal was generated using the Immun-Star AP chemiluminescence kit according to manufacturer’s instructions (BioRad). All blots were imaged using the BioRad ChemiDoc imaging system. Where applicable, relative levels of protein bands were quantified using ImageJ.

### Growth curves

Growth curves were conducted with cultures grown in 96-well plates. Briefly, 2 ml subcultures of all strains were grown for 1 hour at 37°C with shaking at 225 RPM. At 1 hour, 0.1 mM IPTG was added to each culture and 200 µl was aliquoted into 96-well plates. This plate was then grown in a Cytation5 plate reader (BioTek) at 37°C with continuous shaking. OD600 was read every 30 minutes for 24 hours and curves were plotted in Prism (Graphpad).

## Acknowledgements

We would like to thank Dr. Ratmir Derda for the use of the Cytation 5 plate reader; Vincent Man for providing strain VM36 (TR50 Δ*nlpE*) used in this study; and the members of the Tracy Raivio and Jon Dennis labs for many fruitful discussions during the formation of this manuscript.

## Supplemental Materials

**Table S1.** Strains used in this study

| Strain | Description | Source |
| --- | --- | --- |
| MC4100 | F<sub>-</sub> *araD139 (argF-lac)U169 rpsL150 (Strr) relA1 flbB5301 decC1 ptsF25 rbsR* | (1) |
| TR50 | MC4100 λRS88[*cpXP'-lacZ*<sup>+</sup>] | (2) |
| VM36 | TR50 Δ*nlpE* | This study (Vincent Man) |
| TC451 | TR50 Δ*nlpE ompA::kan* | This study |
| TC70 | TR50 + pCA-*nlpE* | This study |
| TC209 | TR50 + pTrc-*nlpE*<sub>WT</sub> | This study |
| TC210 | TR50 + pTrc-*nlpE*<sub>NTD+L</sub> | This study |
| TC211 | TR50 + pTrc-*nlpE*<sub>1-137</sub> | This study |
| TC212 | TR50 + pTrc-*nlpE*<sub>NTD</sub> | This study |
| TC213 | TR50 + pTrc-*nlpE*<sub>1-101</sub> | This study |
| TC214 | TR50 + pTrc99A | This study |
| TC322 | TR50 + pTrc-*nlpE*<sub>CTD+L</sub> | This study |
| TC322 | TR50 + pTrc-*nlpE*<sub>CTD</sub> | This study |
| TC453 | TR50 + pBAD18 | This study |
| TC454 | TR50 + pBAD18-*nlpE*<sub>WT</sub> | This study |
| TC455 | TR50 + pBAD18-*nlpE*<sub>NTD+L</sub> | This study |
| TC468 | TR50 + pBAD18-*nlpE*<sub>DD</sub> | This study |
| TC469 | TR50 + pBAD18-*nlpE*<sub>NTDL+DD</sub> | This study |
| TC272 | TR50 + pCA24N | This study |
| TC273 | TR50 + pCA-*ompA* | This study |
| TC274 | TR50 + pCA-*ompF* | This study |
| TC275 | TR50 + pCA-*ompC* | This study |
| TC276 | TR50 + pCA-*lamB* | This study |
| TC277 | VM36 (TR50 Δ*nlpE*) + pCA24N | This study |
| TC278 | VM36 + pCA-*ompA* | This study |
| TC279 | VM36 + pCA-*ompF* | This study |
| TC280 | VM36 + pCA-*ompC* | This study |
| TC281 | VM36 + pCA-*lamB* | This study |
| TC375 | TR50 + pCA-*ompX* | This study |
| TC376 | TR50 + pCA-*ompW* | This study |
| TC377 | TR50 + pCA-*ompT* | This study |
| TC378 | VM36 + pCA-*ompX* | This study |
| TC379 | VM36 + pCA-*ompW* | This study |
| TC380 | VM36 + pCA-*ompT* | This study |
| TC408 | TR50 + pCA-*ompA*<sub>ΔCTD</sub> | This study |
| TC414 | TR50 + pCA-*ompA*<sub>P198STOP</sub> | This study |
| TC415 | TR50 + pCA-*ompA*<sub>K213A</sub> | This study |
| TC416 | VM36 + pCA-*ompA*<sub>P198STOP</sub> | This study |
| TC417 | VM36 + pCA-*ompA*<sub>K213A</sub> | This study |
| TC462 | TR50 + pCA-*ompA*<sub>PG1</sub> | This study |
| TC463 | TR50 + pCA-*ompA*<sub>PG2</sub> | This study |
| TC464 | TR50 + pCA-*ompA*<sub>PG<sup>-</sup></sub> | This study |
| TC465 | VM36 + pCA-*ompA*<sub>PG1</sub> | This study |
| TC466 | VM36 + pCA-*ompA*<sub>PG2</sub> | This study |
| TC467 | VM36 + pCA-*ompA*<sub>PG<sup>-</sup></sub> | This study |
| TC352 | TR50 Δ*ompA* | This study |
| TC427 | TC352 + pCA24N | This study |
| TC428 | TC352 + pCA-*ompA* | This study |
| TC429 | TC352 + pCA-*ompA*<sub>ΔCTD</sub> | This study |
| TC430 | TC352 + pCA-*ompA*<sub>P198STOP</sub> | This study |
| TC431 | TC352 + pCA-*ompA*<sub>K213A</sub> | This study |
| TC478 | TC352 + pCA-*ompA*<sub>PG1</sub> | This study |
| TC479 | TC352 + pCA-*ompA*<sub>PG2</sub> | This study |
| TC480 | TC352 + pCA-*ompA*<sub>PG<sup>-</sup></sub> | This study |
| TC494 | TC492 + pBAD18-*nlpE*<sub>WT</sub> | This study |
| TC495 | TC492 + pBAD18-*nlpE*<sub>NTD+L</sub> | This study |
| TC472 | OneShot PIR1 + pRE112-*nlpE*<sub>NTD+L</sub> | This study |
| TC493 | MFDpir + pRE112-*nlpE*<sub>NTD+L</sub> | This study |
| TC496 | TR50 *nlpE*<sub>ΔCTD</sub> | This study |
| TC498 | TC496 + pCA24N | This study |
| TC499 | TC496 + pCA-*ompA* | This study |
| TC564 | TC561 + pCA24N | This study |
| TC565 | TC561 + pCA-*ompA* | This study |
| JSW131 | MC4100 Δ*ompA* + pNlp15 (*PcpxP-lux*) | This study |
| JSW158 | MC4100 Δ*cpxA* + pNlp15 | This study |
| JSW160 | MC4100 Δ*nlpE* + pNlp15 | This study |

**Table S2.** Plasmids used in this study

| Plasmid | Description | Source |
| --- | --- | --- |
| pCA24N | Empty ASKA library vector | (3) |
| pCA-*nlpE* | NlpE expression, IPTG-inducible, ASKA library (GFP-) | (3) |
| pCA-*ompA* | OmpA expression, IPTG-inducible, ASKA library (GFP-) | (3) |
| pCA-*ompF* | OmpF expression, IPTG-inducible, ASKA library (GFP-) | (3) |
| pCA-*ompC* | OmpC expression, IPTG-inducible, ASKA library (GFP-) | (3) |
| pCA-*lamB* | LamB expression, IPTG-inducible, ASKA library (GFP-) | (3) |
| pCA-*ompX* | OmpX expression, IPTG-inducible, ASKA library (GFP-) | (3) |
| pCA-*ompW* | OmpW expression, IPTG-inducible, ASKA library (GFP-) | (3) |
| pCA-*ompT* | OmpT expression, IPTG-inducible, ASKA library (GFP-) | (3) |
| pCA-*ompA*<sub>ΔCTD</sub> | Expresses OmpA<sub>Δ194-340</sub>; based on pCA-*ompA* | This study |
| pCA-*ompA*<sub>P198STOP</sub> | Expresses OmpA<sub>1-197</sub>; based on pCA-*ompA* | This study |
| pCA-*ompA*<sub>K213A</sub> | Expresses OmpA<sub>K213A</sub>; based on pCA-*ompA* | This study |
| pCA-*ompA*<sub>PG1</sub> | Expresses OmpA<sub>D262A</sub>; based on pCA-*ompA* | This study |
| pCA-*ompA*<sub>PG2</sub> | Expresses OmpA<sub>R277A</sub>; based on pCA-*ompA* | This study |
| pCA-*ompA*<sub>PG<sup>-</sup></sub> | Expresses OmpA<sub>D262A+R277A</sub>; based on pCA-*ompA* | This study |
| pTrc99A | Empty expression vector, IPTG-inducible expression from *trc* promoter | (4) |
| pTrc-*nlpE*<sub>WT</sub> | NlpE<sub>WT</sub> expression vector; C-terminal 6×His tag | This study |
| pTrc-*nlpE*<sub>NTD+L</sub> | NlpE<sub>NTD+L</sub> expression vector; C-terminal 6×His tag | This study |
| pTrc-*nlpE*<sub>1-137</sub> | NlpE<sub>1-137</sub> expression vector; C-terminal 6×His tag | This study |
| pTrc-*nlpE*<sub>NTD</sub> | NlpE<sub>NTD</sub> expression vector; C-terminal 6×His tag | This study |
| pTrc-*nlpE*<sub>1-101</sub> | NlpE<sub>1-101</sub> expression vector; C-terminal 6×His tag | This study |
| pTrc-*nlpE*<sub>CTD+L</sub> | NlpE<sub>CTD+L</sub> expression vector; C-terminal 6×His tag | This study |
| pTrc-*nlpE*<sub>CTD</sub> | NlpE<sub>CTD</sub> expression vector; C-terminal 6×His tag | This study |
| pBAD18 | Empty expression vector, arabinose-inducible | (5) |
| pBAD18-*nlpE*<sub>WT</sub> | NlpE<sub>WT</sub> expression vector; C-terminal 6×His tag | This study |
| pBAD18-*nlpE*<sub>NTD+L</sub> | NlpE<sub>NTD+L</sub> expression vector; C-terminal 6×His tag | This study |
| pBAD18-*nlpE*<sub>DD</sub> | Full length NlpE with DD(IM) mutation expression vector; C-terminal 6×His tag | This study |
| pBAD18-*nlpE*<sub>NTDL+DD</sub> | NlpE<sub>NTD+L</sub> with DD(IM) mutation expression vector; C-terminal 6×His tag | This study |
| pJW1 | Vector encoding *cpXP-lux* transcriptional reporter | (6) |

**Table S3.**
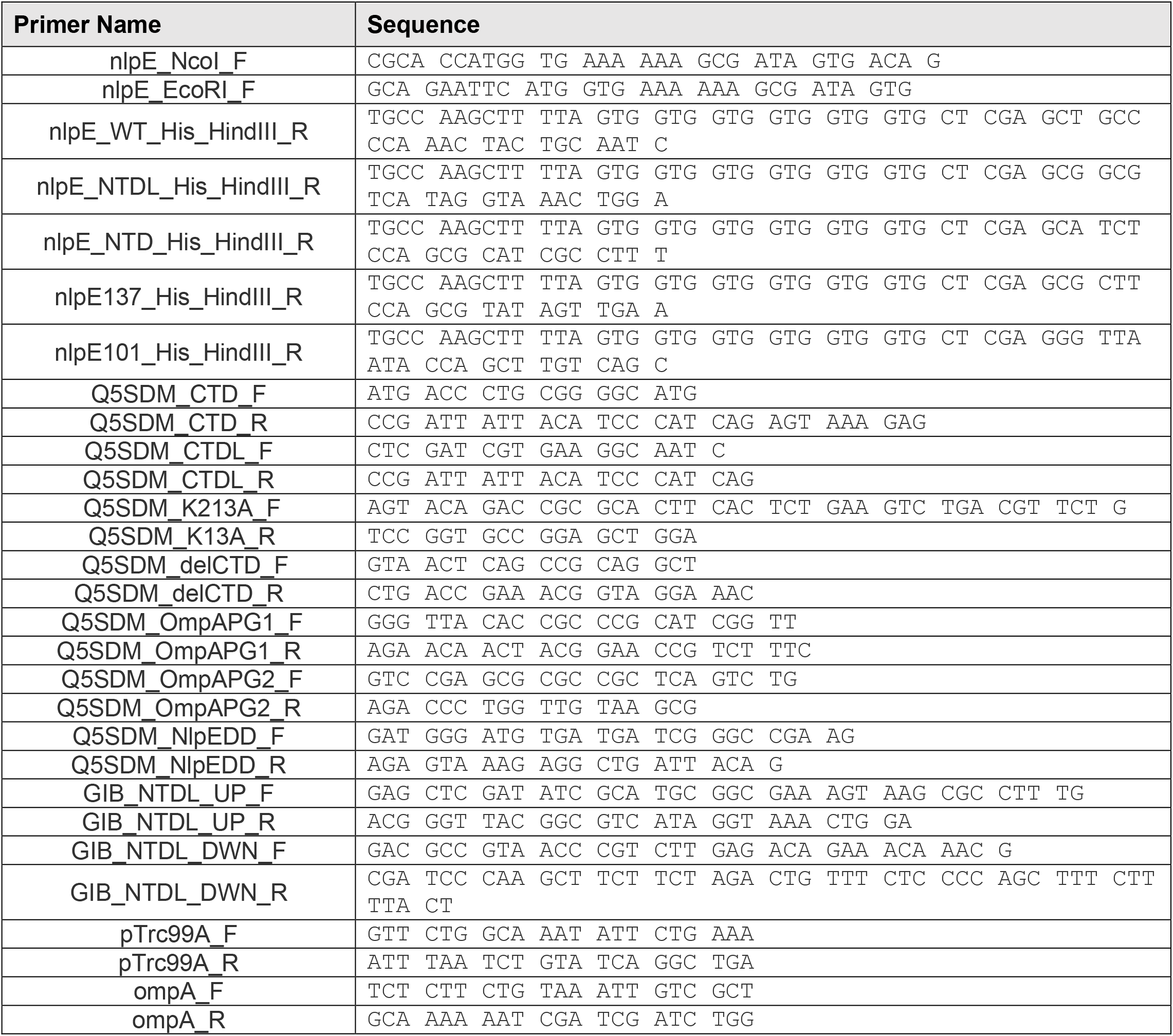
Primers used in this study

**Table S4.** Mass spectrometry results of pull-down assay (30 kDa band)

| Accession | Description | Score<sup>1</sup> | Coverage<sup>2</sup> | # Unique Peptides<sup>3</sup> | # PSMs<sup>4</sup> |
| --- | --- | --- | --- | --- | --- |
| P0A910 | Outer membrane protein A<br>OS=*Escherichia coli* (strain K12)<br>GN=*ompA* PE=1 SV=1 -<br>[OMPA_ECOLI] | 406.33 | 69.08 | 19 | 170 |
| P40710 | Lipoprotein NlpE OS=*Escherichia coli*<br>(strain K12) GN=*nlpE* PE=1 SV=1 -<br>[NLPE_ECOLI] | 294.37 | 78.39 | 16 | 145 |
| C4ZUG9 | 30S ribosomal protein S3<br>OS=*Escherichia coli* (strain K12 /<br>MC4100 / BW2952) GN=*rpsC* PE=3<br>SV=1 - [RS3_ECOBW] | 266.61 | 51.50 | 10 | 100 |

1. Score: Sequest score 2. Coverage: Percent coverage of the protein observed 3. #Unique Peptides: Unique peptides identified that only occur in the protein identified 4. #Peptides: All unique peptide plus peptides that may be common between two or more proteins 5. #PSMs: Number of peptide spectral matches

**Figure S1.**
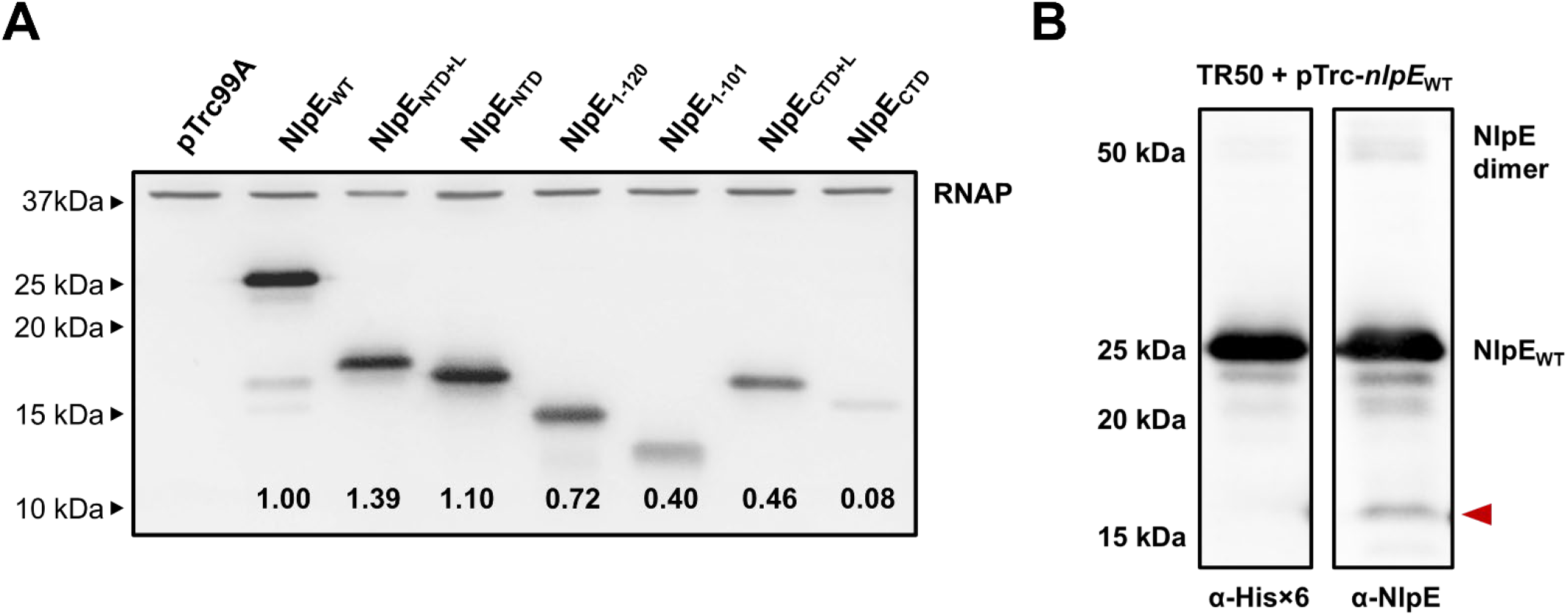
Expression levels of NlpE variants. **(A)** Western blots with anti-His×6 and anti-RNAP (loading control) antibody. Strains containing pTrc99A-based NlpE overexpression vectors were grown to mid-log phase at 37°C with shaking, and NlpE expression was induced with 0.1 mM IPTG for 30 minutes. Cultures were standardized to the same OD, lysed with sample buffer with heating, and subjected to SDS-PAGE and Western blotting analysis. Levels of NlpE were standardized to RNAP loading controls and then standardized to level of WT NlpE. **(B)** Western blot showing NlpE dimerization during overexpression even in denaturing gels as well as a degradation product (red arrowhead) detected by anti-NlpE but not anti-His×6 antibody.

**Figure S2.**
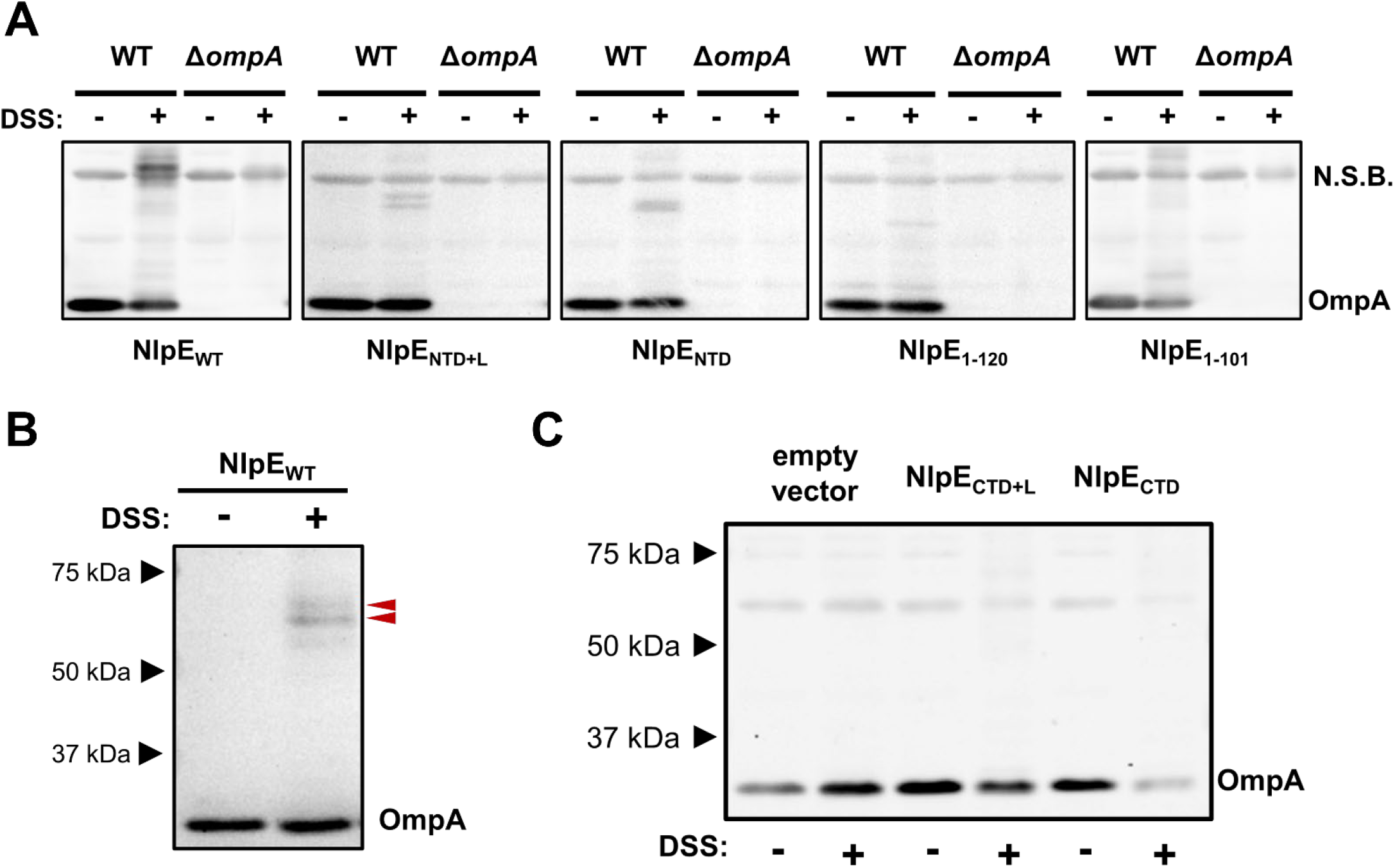
Crosslinking experiments with OmpA. Cells were grown to mid-log (0.5-0.6), induced with 0.1 mM IPTG for 30 minutes, and subjected to crosslinking with 0.5 mM DSS for 30 minutes. Samples were analyzed by SDS-PAGE and Western blotting with anti-OmpA antibody. **(A)** NlpE-OmpA complexes were verified by expressing NlpE in *ompA* deletion background. **(B)** The presence of a double band between NlpE WT and OmpA is visible in the absence of non-specific banding. **(C)** NlpE CTD containing constructs do not crosslink to OmpA under the tested conditions.

**Figure S3.**
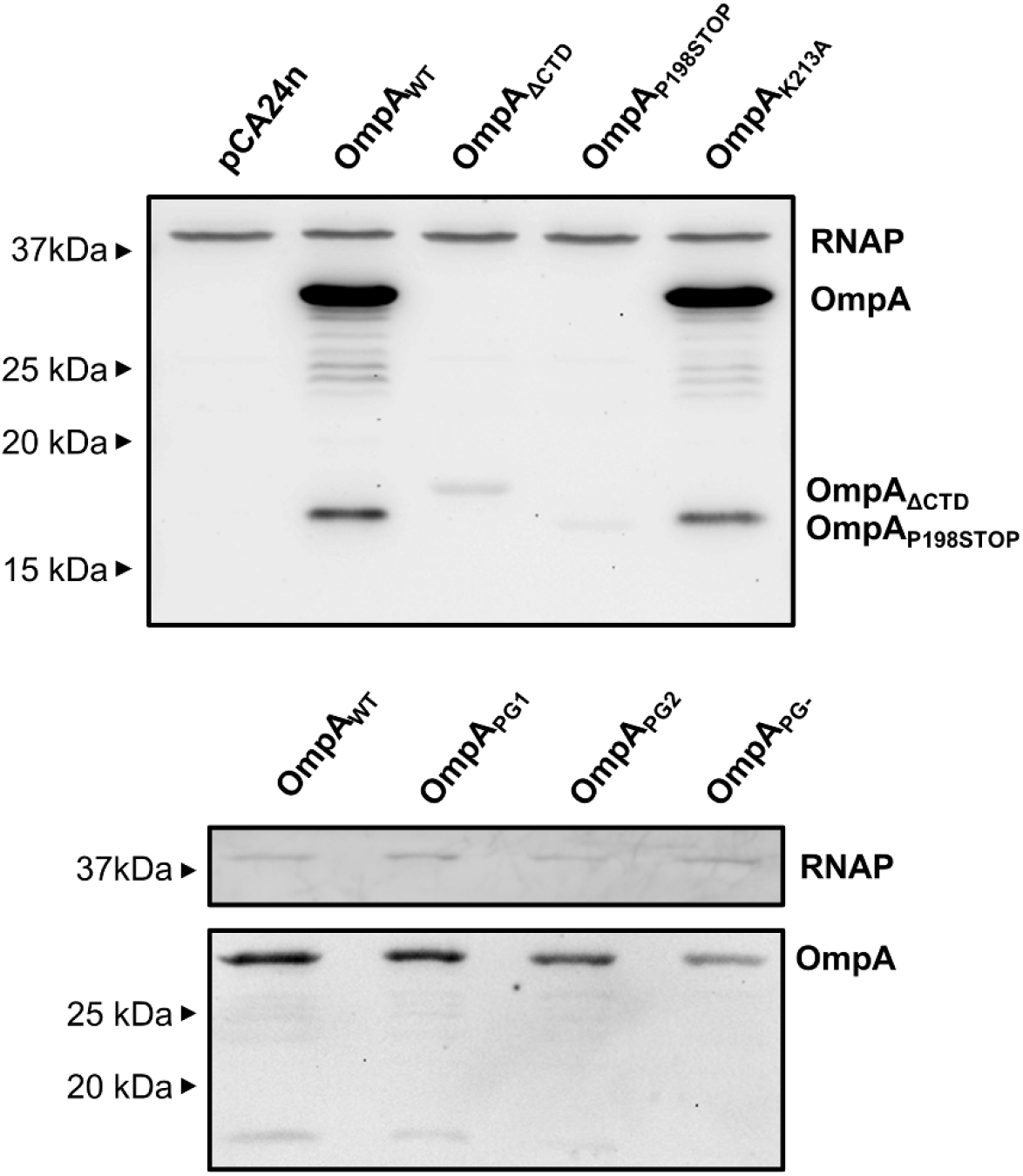
Expression levels of OmpA variants. pCA-*ompA* vectors expressing variants generated by site-directed mutagenesis were transformed into TR50 Δ*ompA* strain. Cultures were grown for 2 hours at 37°C with shaking and induced with 0.1 mM IPTG for 1 hour. An OD-standardized amount of culture was collected, washed, and processed for SDS-PAGE and Western blotting analysis with anti-OmpA antibody. Anti-RNAP antibody was used as a loading control.

**Figure S4.**
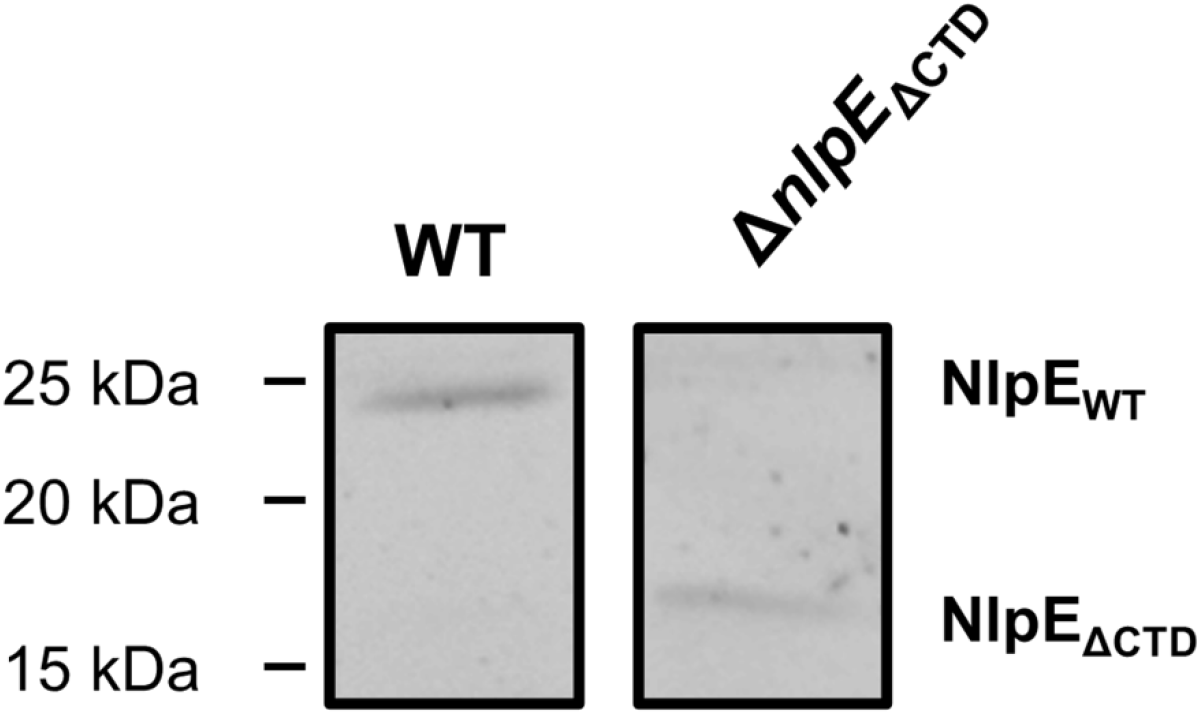
Expression levels of WT NlpE and NlpE_ΔCTD_ expressed from their native chromosomal loci. The immunoblots were conducted on crude membrane fractions isolated by ultracentrifugation.

**Figure S5.**
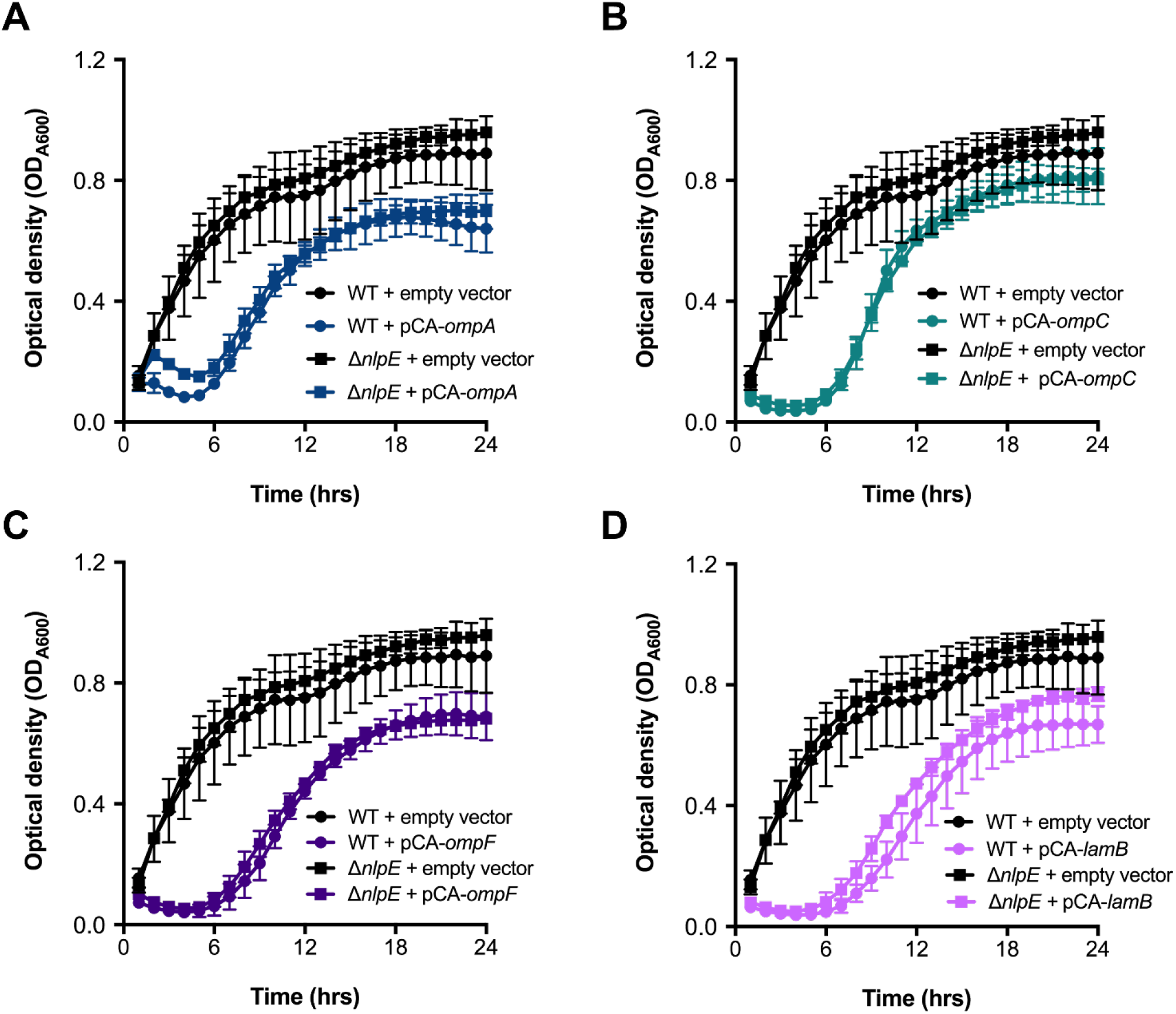
*Growth curves of WT and Δ*nlpE *cells overexpressing OMPs*. WT and Δ*nlpE* TR50 expressing various OMPs were subcultured for 1 hour and induced with 0.1 mM IPTG. Plates were incubated with continuous shaking and OD was read ever 30 minutes for 23.5 in a Cytation5 plate reader. Shown are the means of 4 biological replicates with standard deviations. The OMPS overexpressed were **(A)** OmpA, **(B)** OmpC, **(C)** OmpF, and **(D)** LamB.

## References

1. Yamamoto, K., Hirao, K., Oshima, T., Aiba, H., Utsumi, R., and Ishihama, A. 2005. Functional characterization in vitro of all two-component signal transduction systems from Escherichia coli. J Biol Chem. 280(2):1448–56.

2. Vogt, S.L. and Raivio, T.L. 2012. Just scratching the surface: an expanding view of the Cpx envelope stress response. FEMS Microbiol Lett. 326(1):2–11.

3. Buelow, D.R. and Raivio, T.L. 2005. Cpx signal transduction is influenced by a conserved N-terminal domain in the novel inhibitor CpxP and the periplasmic protease DegP. J Bacteriol. 187(19):6622–30.

4. Isaac, D.D., Pinkner, J.S., Hultgren, S.J., and Silhavy, T.J. 2005. The extracytoplasmic adaptor protein CpxP is degraded with substrate by DegP. Proc Natl Acad Sci U S A. 102(49):17775–9.

5. Snyder, W.B., Davis, L.J.B., Danese, P.N., Cosma, C.L., and Silhavy, T.J. 1995. Overproduction of NlpE, a New Outer Membrane Lipoprotein Suppresses the Toxicity of Periplasmic LacZ by Activation of the Cpx Signal Transduction Pathway. J Bacteriol. 177(15):4216–23.

6. Otto, K. and Silhavy, T.J. 2002. Surface sensing and adhesion of Escherichia coli controlled by the Cpx-signaling pathway. Proc Natl Acad Sci U S A. 99(4):2287–92.

7. Shimizu, T., Ichimura, K., and Noda, M. 2016. The Surface Sensor NlpE of Enterohemorrhagic Escherichia coli Contributes to Regulation of the Type III Secretion System and Flagella by the Cpx Response to Adhesion. Infect Immun. 84(2):537–49.

8. Grabowicz, M. and Silhavy, T.J. 2017. Redefining the essential trafficking pathway for outer membrane lipoproteins. Proc Natl Acad Sci U S A. 114(18):4769–4774.

9. May, K.L., Lehman, K.M., Mitchell, A.M., and Grabowicz, M. 2019. A Stress Response Monitoring Lipoprotein Trafficking to the Outer Membrane. mBio. 10(3).

10. Delhaye, A., Laloux, G., and Collet, J.F. 2019. The Lipoprotein NlpE Is a Cpx Sensor That Serves as a Sentinel for Protein Sorting and Folding Defects in the Escherichia coli Envelope. J Bacteriol. 201(10).

11. Hirano, Y., Hossain, M.M., Takeda, K., Tokuda, H., and Miki, K. 2007. Structural studies of the Cpx pathway activator NlpE on the outer membrane of Escherichia coli. Structure. 15(8):963–76.

12. Orme, R., Douglas, C.W., Rimmer, S., and Webb, M. 2006. Proteomic analysis of Escherichia coli biofilms reveals the overexpression of the outer membrane protein OmpA. Proteomics. 6(15):4269–77.

13. Ortiz-Suarez, M.L., Samsudin, F., Piggot, T.J., Bond, P.J., and Khalid, S. 2016. Full-Length OmpA: Structure, Function, and Membrane Interactions Predicted by Molecular Dynamics Simulations. Biophys J. 111(8):1692–1702.

14. Smith, S.G., Mahon, V., Lambert, M.A., and Fagan, R.P. 2007. A molecular Swiss army knife: OmpA structure, function and expression. FEMS Microbiol Lett. 273(1):1–11.

15. Zakharian, E. and Reusch, R.N. 2003. Outer membrane protein A of Escherichia coli forms temperature-sensitive channels in planar lipid bilayers. FEBS Letters. 555(2):229–235.

16. Zakharian, E. and Reusch, R.N. 2005. Kinetics of folding of Escherichia coli OmpA from narrow to large pore conformation in a planar bilayer. Biochemistry. 44(17):6701–6707.

17. Marcoux, J., Politis, A., Rinehart, D., Marshall, D.P., Wallace, M.I., Tamm, L.K., and Robinson, C.V. 2014. Mass spectrometry defines the C-terminal dimerization domain and enables modeling of the structure of full-length OmpA. Structure. 22(5):781–90.

18. Zheng, C., Yang, L., Hoopmann, M.R., Eng, J.K., Tang, X., Weisbrod, C.R., and Bruce, J.E. 2011. Cross-linking measurements of in vivo protein complex topologies. Mol Cell Proteomics. 10(10):M110 006841.

19. Park, J.S., Lee, W.C., Yeo, K.J., Ryu, K.S., Kumarasiri, M., Hesek, D., Lee, M., Mobashery, S., Song, J.H., Kim, S.I., Lee, J.C., Cheong, C., Jeon, Y.H., and Kim, H.Y. 2012. Mechanism of anchoring of OmpA protein to the cell wall peptidoglycan of the gram-negative bacterial outer membrane. FASEB J. 26(1):219–28.

20. Wang, Y. 2002. The function of OmpA in Escherichia coli. Biochem Biophys Res Commun. 292(2):396–401.

21. Ma, Q. and Wood, T.K. 2009. OmpA influences Escherichia coli biofilm formation by repressing cellulose production through the CpxRA two-component system. Environ Microbiol. 11(10):2735–46.

22. Vogt, S.L. and Raivio, T.L. 2014. Hfq reduces envelope stress by controlling expression of envelope-localized proteins and protein complexes in enteropathogenic Escherichia coli. Mol Microbiol. 92(4):681–97.

23. Konovalova, A., Perlman, D.H., Cowles, C.E., and Silhavy, T.J. 2014. Transmembrane domain of surface-exposed outer membrane lipoprotein RcsF is threaded through the lumen of beta-barrel proteins. Proc Natl Acad Sci U S A. 111(41):E4350–8.

24. Konovalova, A., Mitchell, A.M., and Silhavy, T.J. 2016. A lipoprotein/beta-barrel complex monitors lipopolysaccharide integrity transducing information across the outer membrane. Elife. 5.

25. Cho, S.H., Szewczyk, J., Pesavento, C., Zietek, M., Banzhaf, M., Roszczenko, P., Asmar, A., Laloux, G., Hov, A.K., Leverrier, P., Van der Henst, C., Vertommen, D., Typas, A., and Collet, J.F. 2014. Detecting envelope stress by monitoring beta-barrel assembly. Cell. 159(7):1652–64.

26. Tata, M. and Konovalova, A. 2019. Improper Coordination of BamA and BamD Results in Bam Complex Jamming by a Lipoprotein Substrate. mBio. 10(3).

27. Hart, E.M., Gupta, M., Wuhr, M., and Silhavy, T.J. 2019. The Synthetic Phenotype of DeltabamB DeltabamE Double Mutants Results from a Lethal Jamming of the Bam Complex by the Lipoprotein RcsF. mBio. 10(3).

28. Dekoninck, K., Letoquart, J., Laguri, C., Demange, P., Bevernaegie, R., Simorre, J.P., Dehu, O., Iorga, B.I., Elias, B., Cho, S.H., and Collet, J.F. 2020. Defining the function of OmpA in the Rcs stress response. eLife. 9.

29. Tata, M., Kumar, S., Lach, S.R., Saha, S., Hart, E.M., and Konovalova, A. 2021. High-throughput suppressor screen demonstrates that RcsF monitors outer membrane integrity and not Bam complex function. Proc Natl Acad Sci U S A. 118(32).

30. Rodriguez-Alonso, R., Letoquart, J., Nguyen, V.S., Louis, G., Calabrese, A.N., Iorga, B.I., Radford, S.E., Cho, S.H., Remaut, H., and Collet, J.F. 2020. Structural insight into the formation of lipoprotein-beta-barrel complexes. Nat Chem Biol. 16(9):1019–1025.

31. Arora, A., Rinehart, D., Szabo, G., and Tamm, L.K. 2000. Refolded outer membrane protein A of Escherichia coli forms ion channels with two conductance states in planar lipid bilayers. Journal of Biological Chemistry. 275(3):1594–1600.

32. Singh, S.P., Williams, Y.U., Miller, S., and Nikaido, H. 2003. The C-terminal domain of Salmonella enterica serovar typhimurium OmpA is an immunodominant antigen in mice but appears to be only partially exposed on the bacterial cell surface. Infection and Immunity. 71(7):3937–3946.

33. Samsudin, F., Ortiz-Suarez, M.L., Piggot, T.J., Bond, P.J., and Khalid, S. 2016. OmpA: A Flexible Clamp for Bacterial Cell Wall Attachment. Structure. 24(12):2227–2235.

34. Asmar, A.T., Ferreira, J.L., Cohen, E.J., Cho, S.H., Beeby, M., Hughes, K.T., and Collet, J.F. 2017. Communication across the bacterial cell envelope depends on the size of the periplasm. PLoS Biol. 15(12):e2004303.

35. O’Toole, G.A. and Wong, G.C. 2016. Sensational biofilms: surface sensing in bacteria. Curr Opin Microbiol. 30:139–146.

36. Berne, C., Ellison, C.K., Ducret, A., and Brun, Y.V. 2018. Bacterial adhesion at the single-cell level. Nat Rev Microbiol. 16(10):616–627.

37. Dorel, C., Vidal, O., Prigent-Combaret, C., Vallet, I., and Lejeune, P. 1999. Involvement of the Cpx signal transduction pathway of E. coli in biofilm formation. FEMS Microbiol Lett. 178:169–75.

38. Prigent-Combaret, C., Brombacher, E., Vidal, O., Ambert, A., Lejeune, P., Landini, P., and Dorel, C. 2001. Complex regulatory network controls initial adhesion and biofilm formation in Escherichia coli via regulation of the csgD gene. J Bacteriol. 183(24):7213–23.

39. Jubelin, G., Vianney, A., Beloin, C., Ghigo, J.M., Lazzaroni, J.C., Lejeune, P., and Dorel, C. 2005. CpxR/OmpR interplay regulates curli gene expression in response to osmolarity in Escherichia coli. J Bacteriol. 187(6):2038–49.

40. Dorel, C., Lejeune, P., and Rodrigue, A. 2006. The Cpx system of Escherichia coli, a strategic signaling pathway for confronting adverse conditions and for settling biofilm communities? Res Microbiol. 157(4):306–14.

41. Jones, C.H., Danese, P.N., Pinkner, J.S., Silhavy, T.J., and Hultgren, S.J. 1997. The chaperone-assisted membrane release and folding pathway is sensed by two signal transduction pathways. EMBO J. 16(21):6394–406.

42. Hung, D.L., Raivio, T.L., Jones, C.H., Silhavy, T.J., and Hultgren, S.J. 2001. Cpx signaling pathway monitors biogenesis and affects assembly and expression of P pili. EMBO J. 20(7):1508–18.

43. Raivio, T.L., Leblanc, S.K., and Price, N.L. 2013. The Escherichia coli Cpx envelope stress response regulates genes of diverse function that impact antibiotic resistance and membrane integrity. J Bacteriol. 195(12):2755–67.

44. Lacanna, E., Bigosch, C., Kaever, V., Boehm, A., and Becker, A. 2016. Evidence for Escherichia coli Diguanylate Cyclase DgcZ Interlinking Surface Sensing and Adhesion via Multiple Regulatory Routes. J Bacteriol. 198(18):2524–35.

45. Hernday, A.D., Braaten, B.A., Broitman-Maduro, G., Engelberts, P., and Low, D.A. 2004. Regulation of the pap epigenetic switch by CpxAR: phosphorylated CpxR inhibits transition to the phase ON state by competition with Lrp. Mol Cell. 16(4):537–47.

46. Nevesinjac, A.Z. and Raivio, T.L. 2005. The Cpx envelope stress response affects expression of the type IV bundle-forming pili of enteropathogenic Escherichia coli. J Bacteriol. 187(2):672–86.

47. Vogt, S.L., Nevesinjac, A.Z., Humphries, R.M., Donnenberg, M.S., Armstrong, G.D., and Raivio, T.L. 2010. The Cpx envelope stress response both facilitates and inhibits elaboration of the enteropathogenic Escherichia coli bundle-forming pilus. Mol Microbiol. 76(5):1095–110.

48. Humphries, R.M., Griener, T.P., Vogt, S.L., Mulvey, G.L., Raivio, T., Donnenberg, M.S., Kitov, P.I., Surette, M., and Armstrong, G.D. 2010. N-acetyllactosamine-induced retraction of bundle-forming pili regulates virulence-associated gene expression in enteropathogenic Escherichia coli. Mol Microbiol. 76(5):1111–26.

49. De Wulf, P., Kwon, O., and Lin, E.C.C. 1999. The CpxRA Signal Transduction System of Escherichia coli: Growth-Related Autoactivation and Control of Unanticipated Target Operons. J Bacteriol. 181(21):6772–8.

50. Price, N.L. and Raivio, T.L. 2009. Characterization of the Cpx regulon in Escherichia coli strain MC4100. J Bacteriol. 191(6):1798–815.

51. Confer, A.W. and Ayalew, S. 2013. The OmpA family of proteins: roles in bacterial pathogenesis and immunity. Vet Microbiol. 163(3-4):207–22.

52. Chao, Y. and Vogel, J. 2016. A 3’ UTR-Derived Small RNA Provides the Regulatory Noncoding Arm of the Inner Membrane Stress Response. Mol Cell. 61(3):352–363.

53. Grabowicz, M., Koren, D., and Silhavy, T.J. 2016. The CpxQ sRNA Negatively Regulates Skp To Prevent Mistargeting of beta-Barrel Outer Membrane Proteins into the Cytoplasmic Membrane. mBio. 7(2):e00312–16.

54. Kimkes, T.E.P. and Heinemann, M. 2018. Reassessing the role of the Escherichia coli CpxAR system in sensing surface contact. PLoS One. 13(11):e0207181.

55. Baba, T., Ara, T., Hasegawa, M., Takai, Y., Okumura, Y., Baba, M., Datsenko, K.A., Tomita, M., Wanner, B.L., and Mori, H. 2006. Construction of Escherichia coli K-12 in-frame, single-gene knockout mutants: the Keio collection. Mol Syst Biol. 2:2006 0008.

56. Datsenko, K.A. and Wanner, B.L. 2000. One-step inactivation of chromosomal genes in Escherichia coli K-12 using PCR products. Proc Natl Acad Sci U S A. 97(12):6640–5.

57. Raivio, T.L. and Silhavy, T.J. 1997. Transduction of Envelope Stress in Escherichia coli by the Cpx Two-Component System. J Bacteriol. 179(24):7724–33.

58. Slauch, J.M. and Silhavy, T.J. 1991. cis-Acting ompF Mutations That Result in OmpR-Dependent Constitutive Expression. J Bacteriol. 173(13):4039–48.

59. Miller, J.H., Experiments in molecular genetics. 1972: Cold Spring Harbor Laboratory.

60. Lu, J., Peng, Y., Arutyunov, D., Frost, L.S., and Glover, J.N. 2012. Error-prone PCR mutagenesis reveals functional domains of a bacterial transcriptional activator, TraJ. J Bacteriol. 194(14):3670–7.

61. Macritchie, D.M., Ward, J.D., Nevesinjac, A.Z., and Raivio, T.L. 2008. Activation of the Cpx envelope stress response down-regulates expression of several locus of enterocyte effacement-encoded genes in enteropathogenic Escherichia coli. Infect Immun. 76(4):1465–75.

62. Dunstan, R.A., Hay, I.D., and Lithgow, T., Defining membrane protein localization by isopycnic density gradients. Methods in Molecular Biology. Vol. 1615. 2017: Humana Press Inc. 81–86.

63. Rabilloud, T., Carpentier, G., and Tarroux, P. 1988. Improvement and simplification of low - background silver staining of proteins by using sodium dithionite. ELECTROPHORESIS. 9(6):288–291.

## References

1. Casadaban, M.J. 1976. Transposition and fusion of the lac genes to selected promoters in Escherichia coli using bacteriophage lambda and Mu. Journal of Molecular Biology. 104(3):541–555.

2. Raivio, T.L. and Silhavy, T.J. 1997. Transduction of Envelope Stress in Escherichia coli by the Cpx Two-Component System. J Bacteriol. 179(24):7724–33.

3. Kitagawa, M., Ara, T., Arifuzzaman, M., Ioka-Nakamichi, T., Inamoto, E., Toyonaga, H., and Mori, H. 2005. Complete set of ORF clones of Escherichia coli ASKA library (a complete set of E. coli K-12 ORF archive): unique resources for biological research. DNA Res. 12(5):291–9.

4. Amann, E., Ochs, B., and Abel, K.J. 1988. Tightly regulated tac promoter vectors useful for the expression of unfused and fused proteins in Escherichia coli. Gene. 69(2):301–315.

5. Guzman, L.-M. and Belin, D. 1995. Tight regulation, modulation, and high-level expression by vectors containing the Arabinose PBAD. Journal of Bacteriology. 177(14):4121.

6. Price, N.L. and Raivio, T.L. 2009. Characterization of the Cpx regulon in Escherichia coli strain MC4100. J Bacteriol. 191(6):1798–815.

